# Nucleosome plasticity is a critical element of chromatin liquid–liquid phase separation and multivalent nucleosome interactions

**DOI:** 10.1101/2020.11.23.391599

**Authors:** Stephen E. Farr, Esmae J. Woods, Jerelle A. Joseph, Adiran Garaizar, Rosana Collepardo-Guevara

## Abstract

Liquid–liquid phase separation (LLPS) of chromatin is an important mechanism that helps explain the membrane-less compartmentalization of the nucleus. Because chromatin compaction and LLPS are collective phenomena, linking their modulation to biophysical features of individual nucleosomes is challenging. Here, we develop a novel multiscale chromatin model that integrates atomistic representations, a chemically-specific coarse-grained model, and a minimal model. In tandem, we devise a transferable Debye-length exchange molecular dynamics approach to achieve enhanced sampling of high-resolution chromatin. We find that nucleosome thermal fluctuations become significant at physiological salt concentrations and destabilize the 30-nm fiber. Nucleosome breathing favors stochastic folding of chromatin and promotes LLPS by simultaneously boosting the transient nature and heterogeneity of nucleosome–nucleosome contacts, and the effective nucleosome valency. Our results put forward the intrinsic plasticity of nucleosomes as a key element in the liquid-like behavior of chromatin, and help reconcile longstanding differences between fiber-based and in vivo chromatin models.

## Introduction

The Eukaryotic nucleus is a highly compartmentalized system that achieves its internal organization entirely without the use of membranes^1^. Inside the nucleus, hundreds of millions of DNA base pairs are densely packed into a highly dynamic and heterogeneous structure known as chromatin^2^. The basic building blocks of chromatin are nucleosomes; each composed of approximately 147 base pairs (bp) of DNA wrapped around a histone protein octamer (two copies each of H2A, H2B, H3, and H4)^3,4^. Within chromatin, successive nucleosomes join together by free DNA linker segments of varying lengths—measured in units of nucleosome repeat lengths (NRL = 147 bp + linker DNA length)—and form a 10-nm fiber that resembles ‘beads-on-a-string’^5^. Nucleosomes have strategically evolved charged and contoured surfaces^6^, along with charged and flexible protruding ‘arms’ (histone tails)^7^ that mediate interactions with one another, enabling folding of the 10-nm fiber and the dense packing of DNA inside cells^8,9^.

The structure of chromatin, beyond the 10-nm fiber, remains an intense topic of research and debate^2,10–12^. The traditional textbook view is that 10-nm chromatin folds into a regular and rigid 30-nm solenoid^13^, zigzag^14–16^, or heteromorphic^17^ fiber. However, accumulating new evidence is now shifting the structural paradigm in favor of the ‘liquid-like’ or ‘fluid-like’ model^2,18^, which suggests that 10-nm chromatin fibers condense into an irregular and dynamic polymorphic ensemble^10,19,20^. The term liquid here is used to emphasize a compact chromatin structure that is absent of long-range translational order, and where nucleosomes can flow and relax easily. Several consistent models where chromatin exhibits a disordered organization based on the 10-nm fiber have been proposed recently, including hierarchical looping^21,22^, nucleosome clutches^23^, multiplex higher order folding^24^ and the sea of nucleosomes^25^.

The liquid-like behavior of chromatin is also consistent with its intrinsic heterogeneity in vivo, e.g., the varying DNA sequences and epigenetic marks, the heterogeneous distributions of post-translational histone modifications, the non-uniform NRLs and presence of nucleosome free regions, and the dynamic nucleosome breathing and sliding motions. Many of these factors can independently enable chromatin polymorphism (i.e., folding of 10-nm fibers into irregular loops, hairpins, and bends) by giving rise to a plethora of nucleosome orientations and interactions ^20^–e.g., irregular nucleosome spacing^20,26^, nucleosome depletion^20,26,27^, heterogeneous on/off dyad binding of linker histone proteins to the nucleosome^28,29^, low linker histone concentrations or subtype variations^29^, inhomogenous distributions of post-translational modifications^30^, and the disordered nature of the linker histone protein^31^. In concert, these factors can further amplify or control chromatin polymorphism^30,32^.

In the past three years, the paradigm of a dynamic liquid-like behavior of nucleosomes within cells has gained significant traction due to the realization that chromatin and its associated multivalent biomolecules can undergo liquid–liquid phase separation (LLPS) in vitro and in cells^33–45^. LLPS is now postulated as a mechanism, alongside others ^46^, to explain genome compartmentalization without the use of physical membranes; e.g., the formation of constitutive heterochromatin via the transcriptional repressor HP1^33,34,45^, as well as the emergence of important liquid-like structures such as the nucleoli, Cajal bodies, and nuclear speckles^47^. In addition, the formation of super-enhancer regions has been recently associated with the LLPS of a combination of transcription factors and co-factors, chromatin regulators, non-coding RNAs, and RNA Polymerase II^37–39,42–44^. The emergence of intranuclear phase separation is intricately linked to the complex and crowded biomolecular environment of the cell nucleus^48,49^; i.e, the nucleoplasm is a highly multicomponent mixture of proteins and nucleic acids with varying compositions across different regions^47^. Formation of diverse phase-separated chromatin compartments becomes thermodynamically stable in specific genomic regions when the concentration of key biomolecules, termed scaffolds^50^, surpasses a threshold; such conditions allow scaffolds—normally dissolved in the nucleoplasm—to drive LLPS by minimizing their free energy through the formation of numerous attractive interactions with one another. Therefore, the features that affect binding among nucleosomes, and between nucleosomes and their chromatin-binding proteins (e.g., their chemical makeups and mechanical properties, along with the microenvironments they are exposed to) are expected to be crucial regulators of intranuclear LLPS. In particular, the capacity of biomolecules to interact with at least three others – in other words, their multivalency – is essential because such biomolecules (e.g., proteins, RNA, DNA, and nucleosomes) must form sufficient transient interconnections to compensate for the entropic loss due to the reduced number of microstates upon demixing ^51,52^.

Our work focuses on assessing an important feature of nucleosomes that likely pertains to chromatin LLPS: their notable plasticity. That is, rather than static building blocks—as routinely considered in large-scale chromatin structural models— nucleosomes are highly dynamic and structurally irregular entities^4,53–57^, being better described as a ‘dynamic family of particles’^58^. Nucleosomes in vivo can pack a broad range of DNA base pairs around the histone core (∼100–170 bp), and can also have varied histone compositions and stoichiometries^58^. In addition, thermal fluctuations cause some of the DNA–histone core interactions to spontaneously break and reform from one end of the nucleosome, while the majority of nucleosomal DNA remains wrapped around the histone core^4,56^—this phenomena, known as ‘nucleosome breathing’, is favored at physiological salt concentrations^59,60^. In vivo, the probability of breathing of different nucleosomes across chromatin is likely very diverse too, as it can be sensibly altered by the post-translational modifications each nucleosome carries ^61–63^ and their particular DNA sequences^62,64,65^. Furthermore, combined factors, like the presence of H3K56Ac with DNA sequence changes, can increase the rate of nucleosome unwrapping by one order of magnitude or more^62^. Recruitment of chromatin-binding proteins can also affect the plasticity of nucleosomes. For instance, binding of multiple Swi6 molecules to a nucleosome has been shown to disrupt the DNA–histone bound state, increasing the exposure of the buried histone core to solvent, and crucially, promoting chromatin LLPS^45^.

Nucleosome structural fluctuations provide a transient opportunity for the binding of transcription factors—the proteins that recognize DNA regulatory sequences and assemble protein complexes needed to modulate gene expression—to DNA, and hence, for transcription^66,67^. However, prior to the nucleosome barrier, pioneer and other transcription factors need to surpass the steric hindrance imposed by nucleosome–nucleosome interactions. This seems particularly challenging if we consider the apparent rigidity of nucleosomes within the 30-nm fiber structural models. Therefore, this begs a fundamental question that we target in this work: What are the physical and molecular factors that modulate the accessibility of nucleosomes within compact assemblies? To resolve this conundrum, assessing how the mesoscale properties of chromatin (e.g., density, flexibility, shape, and size) are impacted by the dynamic behavior of nucleosomes is crucial. In this regard, computational modeling of chromatin structure offers an ideal complement to experiments. While polymer chromatin models trained on experimental datasets^68,69^ can help assess if different structural behaviors or physical mechanisms are consistent with the experimental data, mechanistic computational models ^70^ can generate and test new hypotheses based on the physicochemical properties of chromatin. Among these, coarse-grained models with nucleosome and sub-nucleosome resolution^8,9,17,20–22,71–83^ have been particularly useful at providing mechanistic, molecular, and physicochemical understanding on how nucleosome and DNA characteristics impact the mesoscale properties of chromatin.

In the present study, we develop an innovative mechanistic multiscale chromatin model that allows us to investigate the connection between the fine molecular details of nucleosomes (e.g., amino-acid and DNA sequence heterogeneity, specific distributions of post-translational modifications and epigenetic marks, protein flexibility, DNA mechanics, and nucleosome plasticity) and the mesoscale (up to sub-Mb scale) organization of chromatin. By bridging atomistic to sub-Mb chromatin scales, our multiscale approach can dissect the collective behavior emerging from the interactions of a few thousand nucleosomes to propose novel molecular and biophysical mechanism that explain the self-organization and intrinsic LLPS of chromatin. When we model chromatin at residue/base-pair resolution, the dominant role of electrostatics, the high dimensionality of the problem, and the resemblance of chromatin to a poly-branched polymer make sampling highly non-trivial. To overcome this, we develop a powerful Debye-length Hamiltonian replica exchange molecular dynamics (MD) approach that is transferable and can be used to explore the energy landscapes of other intractable charged systems.

Consistent with experiments^59,60^, our multiscale model and Debye-length sampling technique reveal that nucleosome breathing is enhanced at physiological salt conditions. In turn, we observe that while strengthening of nucleosome breathing drives chromatin to populate a highly dynamical, but compact, liquid-like structural ensemble^2,18^, inhibition of breathing gives rise to 30-nm regular fibers. Our simulations further explain that liquid-like chromatin organization is characterized by short-lived and orientationally-diverse internucleosome interactions mediated by transient non-specific DNA–histone tail contacts, whereas 30-nm fibers are sustained by long-lived regular face-to-face nucleosome interactions. Importantly, nucleosome plasticity promotes both liquid-like folding of individual chromatin systems and LLPS of chromatin arrays via the same physical mechanism: namely, it enhances the multivalency of nucleosomes and, therefore, the connectivity and stability of both compact chromatin and the phase-separated condensed chromatin liquid^52^. The stochastic organization of nucleosomes within compact chromatin that we observe, both within single arrays and phase-separated liquid drop compartments, paints a much more permissive picture of nucleosome targeting than that offered by the fiber-based models. The realization that nucleosomes can be simultaneously stochastically and tightly packed might have important implications in reshaping the molecular mechanisms used to link chromatin structure to modulation of DNA accessibility.

## Results

### Multiscale model for heterogeneous chromatin

We have designed our multiscale model (Figures 1 and S1) to capture the physicochemical heterogeneity of chromatin and the plasticity of nucleosomes observed in vivo. To describe these features on functionally relevant length scales (i.e., up to sub-Mb scales), we integrate three complementary levels of resolution: atomistic representations of nucleosomes, a chemically-specific chromatin model, and a minimal chromatin model (the detailed description of our models is given in ‘Methods’ and the Supporting Information). Our mid-resolution layer is termed our ‘chemically-specific coarse-grained model’ as it features representations of breathing nucleosomes with all histone proteins resolved at the residue level (preserving their charge, hydrophobicity, size, and atomistic flexibility) and double-stranded DNA at the base-pair step level (with charge and sequence-dependent mechanical properties described with a modified version of the Rigid Base Pair (RBP)^84–88^ model with added phosphate charges; see ‘Methods’). Parameters for our chemically-specific model are obtained from experimental amino-acid pairwise contact propensities^89–91^, large datasets of atomistic MD simulations of DNA strands^92^, and bias-exchange metadynamics atomistic simulations of 211-bp nucleosomes^31^. Notably, the physicochemical and molecular fidelity of histones and DNA within our chemically-specific coarse-grained model give rise to a DNA polymer that spontaneously wraps ∼1.7 times around the histone core, and that adopts the correct topology and left-handed chirality under weak negative DNA supercoiling, and the chiral inverted right-handed counterpart under weak positive supercoiling, consistent with experiments^93–96^ (see Supporting Information and Figure S6). The molecular resolution of our model also results in nucleosomes that naturally exhibit spontaneous breathing motions (i.e., without being primed to do so) and display force-induced unwrapping in quantitative agreement with experiments at single base-pair resolution (see ‘Model Validation’, Supporting Information, and Figure 2).

**Figure 1.**
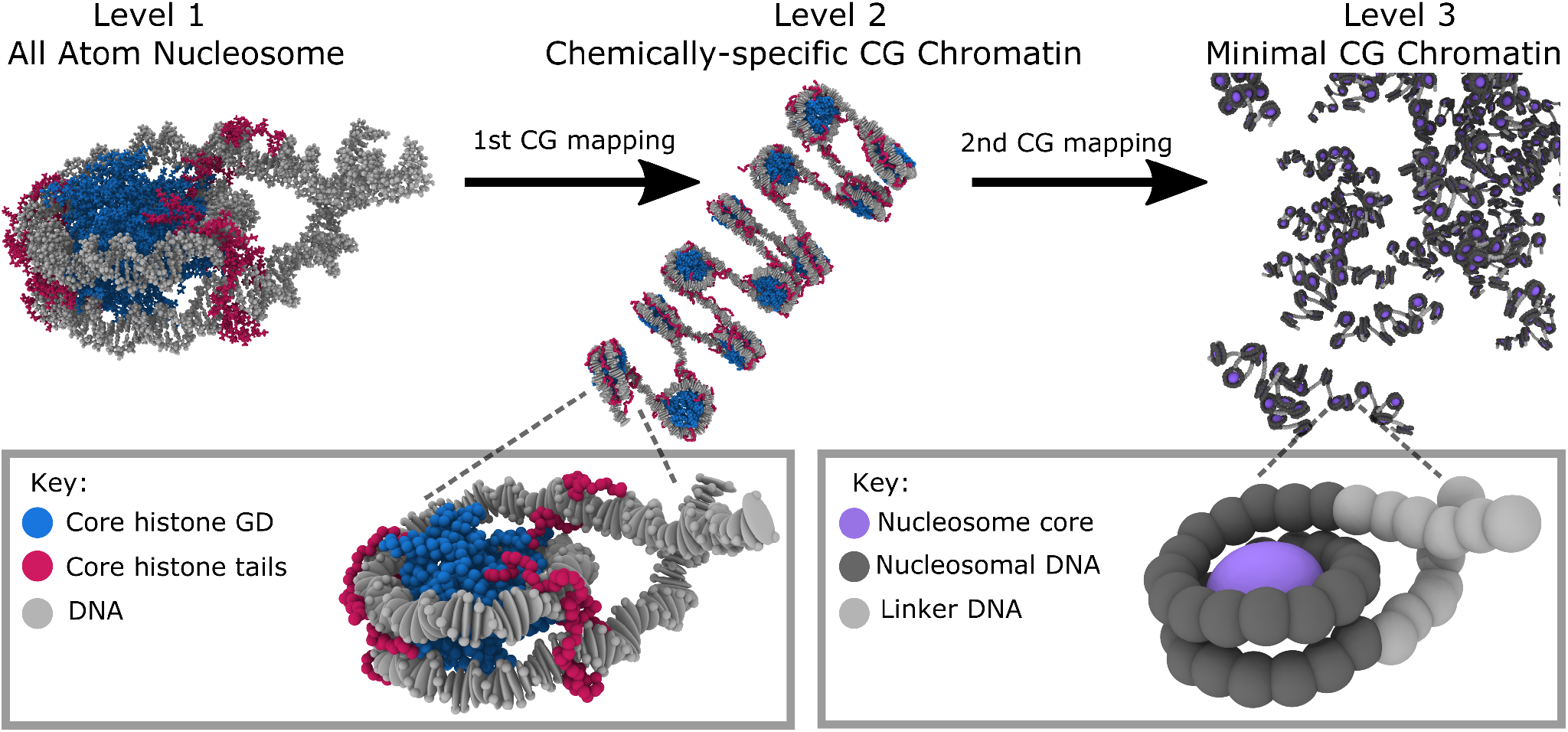
Schematic representation of our chromatin multiscale toolkit spanning three levels of resolution. **(Level 1)** Atomistic MD simulations of DNA and nucleosomes to identify key physicochemical information. **(Level 2)** Debye-length replica exchange MD simulations of our chemically-specific coarse-grained chromatin model; representing DNA at the base-pair level (one grey elipsoid per base-pair) and histone tails (one magenta bead per amino acid) and globular domains (GD, one blue bead per amino acid) at the residue level. This model is able to link elementary properties of nucleosomes to mesoscale behavior of oligonucleosomes as it accounts for the following key molecular features from Level 1: DNA mechanical properties, secondary structure of the histone globular regions, flexibility of histone tails, and the size, shape, electrostatics and hydrophobicity of individual amino acids and base-pairs. **(Level 3)** Direct coexistence simulations of our minimal coarse-grained chromatin model designed to investigate the phase behavior of a few thousand interacting nucleosomes. The histone core is treated as single interaction site (purple bead) with parameters from internucleosome potential of mean force simulations from Level 2. The nucleosomal DNA (dark grey beads) and the linker DNA (light grey beads) are described explicitly (1 bead = 5 bp) with our minimal RBP-like model parameterized from free DNA simulations from Level 2.

**Figure 2.**
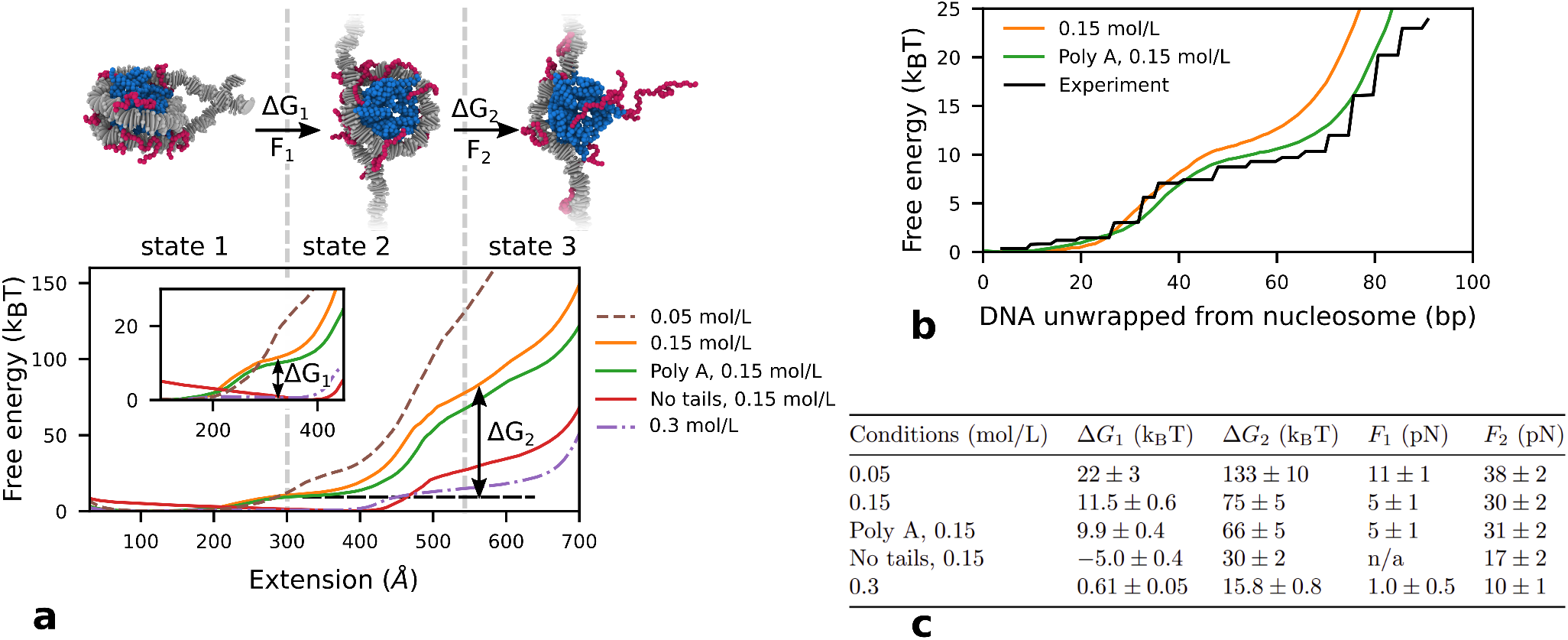
Validation of chemically-specific coarse-grained model against force spectroscopy experiments. **a** Model predictions for the force-induced unwrapping of mononucleosomes under varying conditions. Top: Representative simulation snapshots of nucleosome configurations (color coded as in Figure 1–Level 2) at three different stages of the unwrapping process, showing a fully wrapped nucleosome (state 1) at low pulling forces (≤*F*_1_ in Table in **c**), a nucleosome with the first turn unwrapped (state 2) at intermediate forces (*F*_1_–*F*_2_ in Table in **c**), and a fully unwrapped nucleosome (state 3) at higher forces (≥ *F*_2_ in Table in **c**). Bottom: Free energy cost for nucleosome unwrapping as measured by the PMF as a function of the end-to-end DNA distance (or extension). The dashed brown, solid orange, and dashed purple curves correspond to 601-nucleosome simulations at 0.05 mol/L, 0.15 mol/L, and 0.3 mol/L of NaCl, respectively. The green curve corresponds to simulations of a poly-A nucleosome at 0.15 mol/L NaCl. The red curve was calculated for a nucleosome with all histone tails clipped at 0.15 mol/L NaCl. The inset provides a zoomed in view of the low-force regime and indicates the free energy difference between states 1 and 2 (Δ*G*_1_) for the simulations of 601-nucleosomes at 0.15 mol/L NaCl. The vertical dashed lines are used as visual aids to guide the reader to the approximate regions on the PMF that exhibit different states of nucleosome unwrapping. The horizontal dashed line highlights the free energy plateau corresponding to the transition between states 1 and 2 of the 601-nucleosomes at 0.15 mol/L NaCl; the free energy difference between states 2 and 3 (Δ*G*_2_) is also illustrated for this case. **b** Quantitative agreement between the free energy of nucleosome unwrapping at single DNA base-pair resolution estimated with our simulations at 0.15 mol/L NaCl for nucleosomes with the 601 sequence (orange curve) and a poly-A sequence (green curve), and that derived from analysis of mechanical unzipping experiments (black curve)^105,106^ **c** Summary of the change in free energy (mean *±* standard deviation) between nucleosome unwrapping states, and the corresponding rupture forces. Δ*G*_1_ is the free energy difference between states 1 and 2. Δ*G*_2_ is the free energy difference between states 2 and 3. *F*_1_ is the maximum force during the state 1 to state 2 transition, *F*_2_ is the maximum force during the state 2 to state 3 transition. n/a for no tails denotes that there is no rupture force for state 1 to state 2 transition as the free energy minimum is state 2. The values of *G* and *F* are obtained from reading off the graphs in **a** and the uncertainty values come from the range of positions that these values could be read from.

Further coarse-graining is essential to reduce the system dimensionality and investigate chromatin LLPS. Accordingly, from our chemically-specific coarse-grained simulations, we derive a consistent minimal chromatin model that describes each nucleosome with just a few particles and enables the simulation of chemically heterogeneous sub-Mb scale chromatin regions and LLPS, while also considering nucleosome thermal fluctuations. Specifically, we use one ellipsoid for each histone core, and develop a ‘minimal RBP-like model’ for DNA at a resolution of 5-base-pairs per bead, with minimal helical parameters extracted from chemically-specific coarse-grained simulations of 200 bp DNA strands (see ‘Methods’ and Supporting Information). Nucleosome–nucleosome and nucleosome–DNA interactions are modeled with orientationally-dependent potentials fitted to reproduce internucleosome potentials of mean force calculated with our chemically-specific coarse-grained chromatin model (see Figure S4 and Supporting Information). Because our multiscale approach preserves amino-acid and DNA sequence, it can be used to investigate large-scale chromatin structural implications of subtle chemical changes (e.g., in charge, hydrophobicity and flexibility) that might stem from protein/DNA mutations, post-translational modifications of histones, or DNA epigenetic marks.

### Model validation

We begin by comparing various quantities in our chemically-specific coarse grained chromatin model with corresponding experimental observables. First, we find that the persistence length of DNA estimated in our simulations (i.e., performed using our modified RBP model with phosphate charges; see ‘Methods’) agrees well with salt-dependent values from high-throughput tethered particle motion single-molecule^97^ and light scattering^98^ experiments, and with sequence-dependent measurements from cyclization assays^99^ (see Figure S7 and Supporting Information). The residue-resolution protein model^89^ that we use to describe histones has been shown to reproduce well the experimental radii of gyration of many intrinsically disordered proteins^91^.

Next, we assess the accuracy of the critical dynamical unwrapping behavior of nucleosomes in our model by performing single-molecule force-extension simulations and comparing our results with those from force spectroscopy experiments^100–103^. Pulling a single nucleosome at the equilibrium speeds typically used in force spectroscopy in vitro experiments (e.g., ~0.1 mm/s for magnetic tweezers^103^) is computationally unfeasible, as this would require millisecond-long trajectories, which are presently not easily achievable. With the current computing power, force-extension steered MD would need to be performed at pulling speeds significantly above those required to maintain equilibrium conditions (e.g., 10–100 times faster than in experiments). To overcome this computational limitation, following the procedure of Lequieu et al.^104^, we perform equilibrium umbrella sampling to estimate the potential of mean force (PMF) of mononucleosome unwrapping, using the DNA end-to-end distance as the order parameter. Subsequently, we derive force-extension plots by taking the numerical derivative of the PMF curves (see force-extension curves in Figure S8).

Consistent with experiments, the force-induced nucleosome unwrapping behavior predicted by our model can be separated into three equilibrium regimes (Figure 2a,b), each spanning a force-extension region (Figure S8) that matches the experimental values in Refs. 100–103 (see further discussion in Supporting Information). Most excitingly, by modeling nucleosomes with chemical and mechanical accuracy, we obtain quantitative agreement between our model predictions and the free energy landscape for nucleosome unwrapping^105^ derived from mechanical unzipping experiments at a single DNA base-pair resolution^106^ (Figure 2c).

Further, our model shows that unwrapping of the outer DNA turn is associated with a free energy barrier (Δ*G*_1_ in Figure 2) of ∼11.5 *k*_*B*_*T* at 0.15 mol/L NaCl, which closely matches estimates from force spectroscopy experiments at similar salt conditions (∼9–11.1 *k*_*B*_*T*)^101,103,107^. In agreement with magnetic tweezer experiments^107^, we estimate an increase of ∼10.5 *k*_*B*_*T* in the free energy barrier for outer DNA turn unwrapping as the monovalent salt concentration decreases from 0.15 mol/L to 0.05 mol/L of NaCl (Figure 2a,c). This increase in free energy can be explained from the increased electrostatic attraction between DNA and histones at low salt. Hence, as previously shown, nucleosome breathing is feasible at physiological salt concentrations, but becomes increasingly challenging in low salt conditions^59^.

The free energy landscape for nucleosome unwrapping computed with our model agrees well with several other experimental findings. For instance, in line with the dependency of nucleosome unwrapping on DNA sequence^62^, nucleosomes with an unfavorable polyA DNA sequence have a free energy barrier for unwrapping the outer DNA turn that is 1.5 *k*_*B*_*T* smaller than in 601 nucleosomes. In addition, histone tail clipping leads to unwrapping of the outer turn with negligible energetic penalty, in agreement with mechanical disruption experiments where unwrapping of the outer DNA turn occurs at near-zero forces after histone tail removal^100^.

Regarding chromatin behavior, our force-extension curves computed using an extension clamp (Figure S8) exhibit the typical saw-toothed pattern of optical tweezer experimental curves^108^, where the force displays an abrupt drop accompanied by an increase in the extension due to the partial unwrapping of individual nucleosomes (see further discussion in the Supporting Information). Such behavior is also consistent with the step-like patterns emerging from magnetic tweezer experiments^103^, where a force clamp is used instead.

### Nucleosome plasticity underlies the stochastic folding of chromatin

Our chemically-specific coarse-grained chromatin model contains sufficient physical details to examine the effects of nucleosome breathing on the structure of small (*<*10 kb) chromatin systems. However, simulating chromatin arrays at high resolution (i.e., one bead per protein residue or DNA base pair) in physiological conditions is computationally challenging for various reasons. When represented at high-resolution, chromatin is a high-dimensional system made up of a large number of opposite charged particles (e.g., lysine and DNA phosphate beads) that establish strong long-range interactions with each other, and, hence, the energy landscape of chromatin is populated by many competing low-lying minima separated by high energy barriers. Such a rugged energy landscape is difficult to sample with standard MD simulations, as transitions across the high energy barriers are rare within the accessible simulation timescales. Although Monte Carlo (MC) simulations are effective at overcoming high energy barriers, chromatin at high-resolution and under physiological conditions resembles a highly dense poly-branched polymer; consequently, most MC moves have a low acceptance probability due to steric clashes even after small displacements, just like in a dense liquid. To overcome these challenges and achieve sufficient sampling of chromatin at high resolution, we developed a novel Hamiltonian replica exchange MD scheme that varies the Debye length across replicas. An advantage of this approach, over standard temperature replica-exchange MD (REMD), is that it allows us to use a much smaller number of replicas (i.e., 16 instead of 80 for a 12-nucleosome system). This is achieved by implementing larger differences within the Hamiltonians of neighboring replicas but focused on selected degrees of freedom; such degrees of freedom are chosen to precisely modulate the electrostatic interactions, which contribute most strongly to the high energy barriers (see Supporting Information). In addition, unlike in classical REMD where most high temperature replicas are not physically meaningful, in our approach each of the Debye-length-tuned replicas explores the behavior of chromatin at an independent monovalent salt concentration that is accessible to different experiments (e.g., within the 0.01–0.15 mol/L NaCl range).

Using this Debye-length exchange approach to sample our chemically-specific model, we can compare the behavior of chromatin with nucleosomes that exhibit spontaneous DNA unwrapping (i.e., DNA breathing) versus cases where the nucleosomes are constrained to remain permanently wrapped (i.e., no DNA breathing is allowed). As a benchmark for our model, we focus on 12-nucleosome chromatin arrays with a regular NRL of 165 bp (or a linker length of 18 bp); since in vitro sedimentation coefficients are available for validation^109^, and because short regular NRLs favor folding into ideal zigzag structures well-characterized by near-atomic resolution in vitro experiments^14,15^. Furthermore, using a regular NRL across the array allows us to exclude the structural heterogeneity stemming from linker DNA variability^20^, and to isolate the effects of nucleosome thermal breathing.

Our simulations reveal that constraining the nucleosomal DNA to remain fully and permanently attached to the histone core directs chromatin with short DNA linkers to fold into the 30-nm rigid ladder-like zigzag fibers at 0.15 mol/L NaCl (see ‘Non-breathing’ in Figure 3a); i.e., where nucleosomes stack perfectly face-to-face with their second-nearest neighbors and rarely interact with other nucleosomes or in alternate orientations (Top panel in Figure 3b). The regular zigzag fiber structure we observe is analogous to that derived from cryo-EM experiments^15^ for 12-nucleosome chromatin systems with short linker lengths, and the 167-bp tetranucleosome crystals^14^.

**Figure 3.**
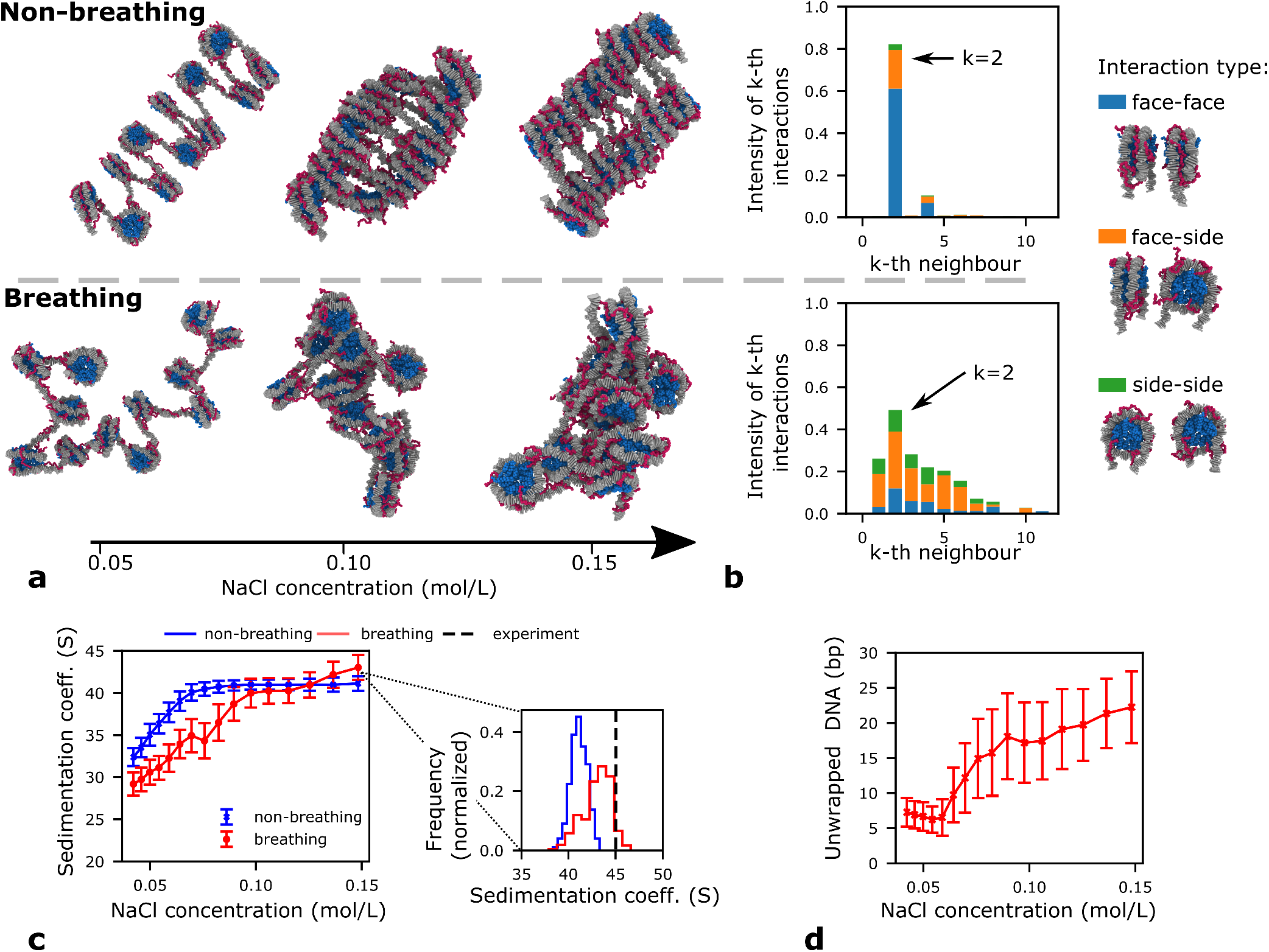
Structural differences in chromatin arrays with breathing versus non-breathing nucleosomes. **a** Representative simulation snapshots of 165-bp 12-nucleosome chromatin with non-breathing (top) versus breathing (bottom) nucleosomes at three different salt concentrations: 0.05 mol/L, 0.10 mol/L and 0.15 mol/L of NaCl. **b** Bar plots depicting the frequency of interactions among k-th nearest nucleosomes neighbors for chromatin with non-breathing (top) versus breathing (bottom) nucleosomes. The bars are colored according to the percentage of the nucleosome pairs that engage in face-to-face (blue), face-to-side (orange), or side-to-side (green) interactions; these types of interactions are illustrating by the cartoons on the right. The definition of the nucleosome axes used to determine if an interaction occurs face-to-face, face-to-side, or side-to-side are given in Figure S9. **c** Sedimentation coefficients versus NaCl concentration (right) for chromatin with non-breathing (blue) versus breathing (red) nucleosomes. Histograms comparing the distributions of sedimentation coefficient values for chromatin with non-breathing (blue solid) and breathing (red solid) at 0.15 mol/L in our simulations with the experimental value from reference^109^ (black dashed). **d** Number of average DNA bp that unwrap per nucleosome in our simulations at varying concentration of NaCl.

More strikingly, our simulations show that the thermal breathing motion of nucleosomes destabilizes the formation of regular 30-nm zigzag fibers, and favors instead the organization of chromatin into a liquid-like ensemble at 0.15 mol/L NaCl (‘Breathing’ in Figure 3a). This liquid-like ensemble encompasses a wide-range of compact structures where nucleosomes interact with a multiplicity of neighbors in diverse orientations (i.e., face-to-face, side-to-side, and face-to-side) (Bottom panel in Figure 3b), as had been postulated by Maeshima and collaborators^19^. Importantly, within this liquid-like ensemble chromatin is characterized simultaneously by a higher degree of flexibility and compaction than chromatin within 30-nm fibers (‘Non-breathing’ in Figure 3c). These features are evident from the wider distributions of the sedimentation coefficients (shifted towards the right), compared to those of chromatin with non-breathing and static nucleosomes, and a mean value at 0.15 mol/L NaCl that is in excellent agreement with the experimental measurements (Figure 3c). Our simulations also capture qualitatively the progressive decondensation of chromatin with decreasing monovalent salt observed experimentally^109^.

To rationalize the impact of nucleosome breathing in chromatin self-assembly, we quantify the number of DNA base pairs that unwrap per nucleosome due to thermal breathing in a given time-frame (Figure 3d). Consistent with in vitro single-molecule fluorescence resonance energy transfer experiments—where nucleosomes are considerably unwrapped at 0.1 mol/L NaCl (i.e., ∼30% probability; 1.5 ms unwrapped versus 3–4 ms wrapped) but minimally unwrapped when salt is lowered to 0.02 mol/L (i.e., ∼10% probability; 1.4 ms unwrapped versus 14 ms wrapped)^59^—we observe that the average number of unwrapped DNA base pairs is most significant at physiological salt (i.e., 22 bp*±*5 bp at high salt vs 7 bp*±*2 bp at low salt); this occurs because the DNA–histone core electrostatic attraction weakens under conditions of high electrostatic screening. Therefore, at physiological salt, the DNA linker regions transiently lengthen and shorten back (i.e., the average length of the linker DNA doubles from 18 to∼38 bp), subsequently increasing the size of the DNA bending fluctuations and the accessible range of internucleosome rotational angles. It is precisely the emergence of these large fluctuations that explains, from a molecular point of view, the origin of the nucleosome–nucleosome orientation heterogeneity that sustains the liquid-like behavior of chromatin. Consistently, mesoscale simulations showed that large DNA linker variations increase the structural heterogeneity of chromatin^20^, and experiments have shown that internucleosome rotational angle variability favors chromatin structural heterogeneity^24,109^. Furthermore, the structural effects of nucleosome breathing that we observe are more pronounced in chromatin arrays with short NRLs, as the internucleosome orientations within such systems are otherwise highly restricted by their short linkers^109^ (Figure S10 and Supporting Information).

This modulation of nucleosome breathing with salt and its impact in chromatin structure may help explain why ordered and disordered structural chromatin models have been derived from in vitro and in vivo data, respectively. Such differences have already been attributed to the low salt concentrations used in many of the in vitro experiments^2,18^, and to the regularity of reconstituted chromatin arrays—i.e., with strong nucleosome positioning sequences and homogeneous linker DNA sequences, uniform NRLs, homogeneous histone protein compositions, and a relatively small number of nucleosomes (∼4–100)^20^. Our work further suggests that considering the disparities in dynamic behavior of nucleosomes at physiological versus low salt is important in such debate.

### Liquid-like chromatin is stabilized by short-lived non-specific DNA–histone tail electrostatic interactions

The ability of our chemically-specific model to resolve the motions of individual amino acids and DNA base pairs within compact chromatin enables us to examine the precise contributions of each of these species in directing chromatin organization. Specifically, we compute the fraction of time each amino acid and DNA base pair in a given nucleosome mediate inter-nucleosome interactions. These interactions are categorized into the three main groups: DNA, globular regions, and histone tails. This analysis reveals that the molecular driving forces that stabilize the liquid-like organization of chromatin and the regular 30-nm fiber folding are strikingly different (Figure 4).

**Figure 4.**
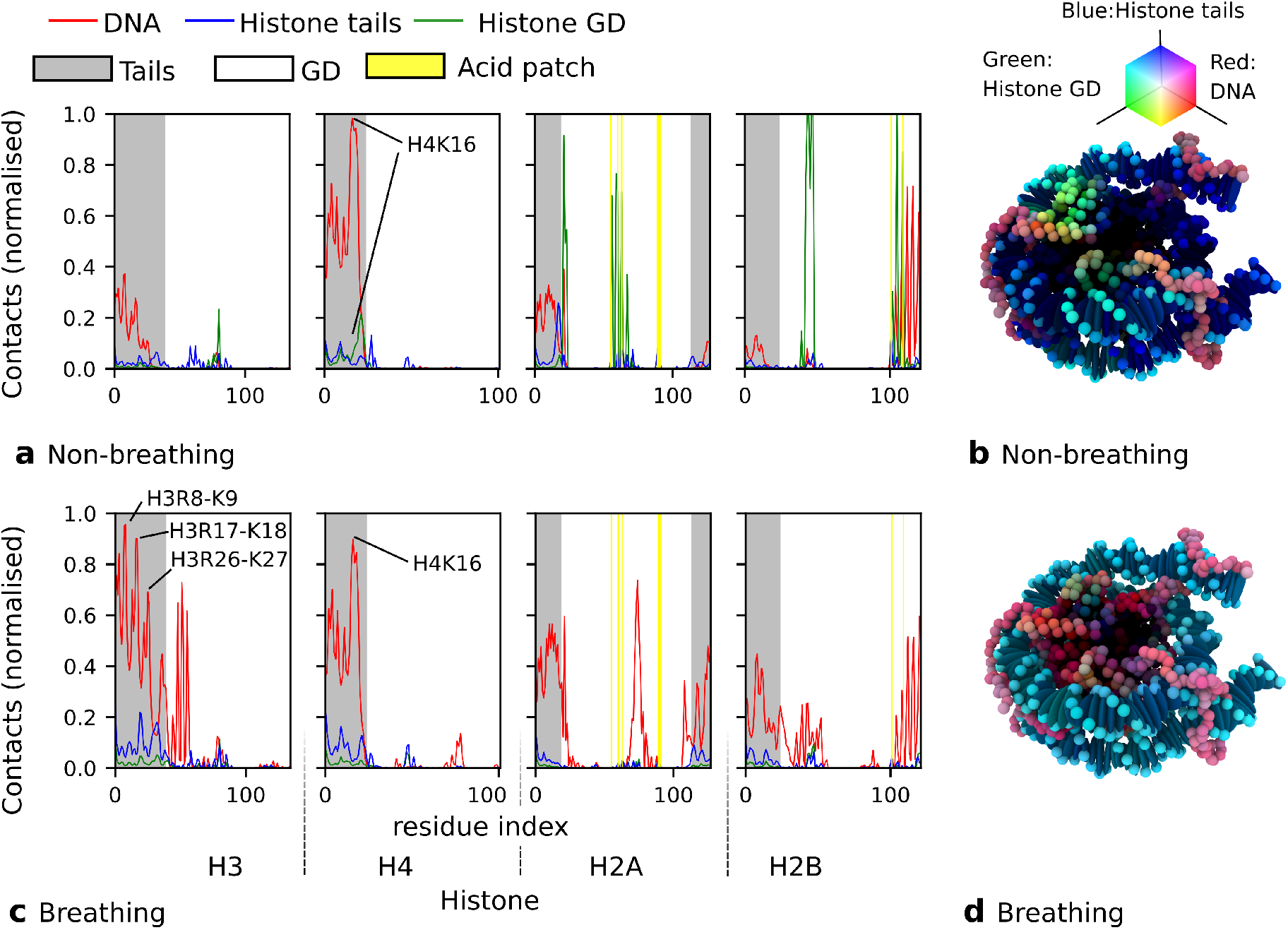
Molecular interactions that sustain chromatin compaction. Normalized fraction of nucleosome–nucleosome interactions (contacts) within compact chromatin ((**a**) with breathing nucleosomes, (**c**) with non-breathing nucleosomes) that are mediated by histone–histone or histone–DNA interactions. The horizontal axis runs across all the histone protein residues within the core (i.e., H3, H4, H2A, and H2B) of a reference nucleosome, with the grey shaded area highlighting its histone tail residues, the white area its globular domain residues, and the yellow vertical lines the residues within the acidic patch. The fraction of contacts are broken down by type: DNA–histone (red), histone tail–histone (blue), and globular domain–histone (green). (**b**,**d**) Visualization of the preferential types of interaction per residue or base pair in non-breathing versus breathing nucleosomes. Each residue and DNA base pair are colored according to the RGB value that is obtained by combining the red, green, and blue values of lines in (**a**,**c**).

In agreement with electron microscopy and single-molecule force spectroscopy experiments on H4-tail cross-linked chromatin^110,111^ showing that 30-nm zigzag fiber folding is stabilized by face-to-face interactions between the H4-tail of one nucleosome and the H2A histone on the surface of another, we observe that 30-nm fibers are sustained by interactions between globular histones on the surfaces of the two stacked nucleosomes (including those within the acidic patch), between the H4 tail and DNA, and more modestly between the H4 tail and the acidic patch (Figure 4a,b). Besides the H4-tail—which has an ideal location on the nucleosome face—the other histone tails play a less prominent role in the folding of the 30-nm fiber. Our simulations further illuminate that the face-to-face stacking of nucleosomes within zigzag fibers is long lived (i.e., once stacked, nucleosomes rarely unstack; Figure 5a).

**Figure 5.**
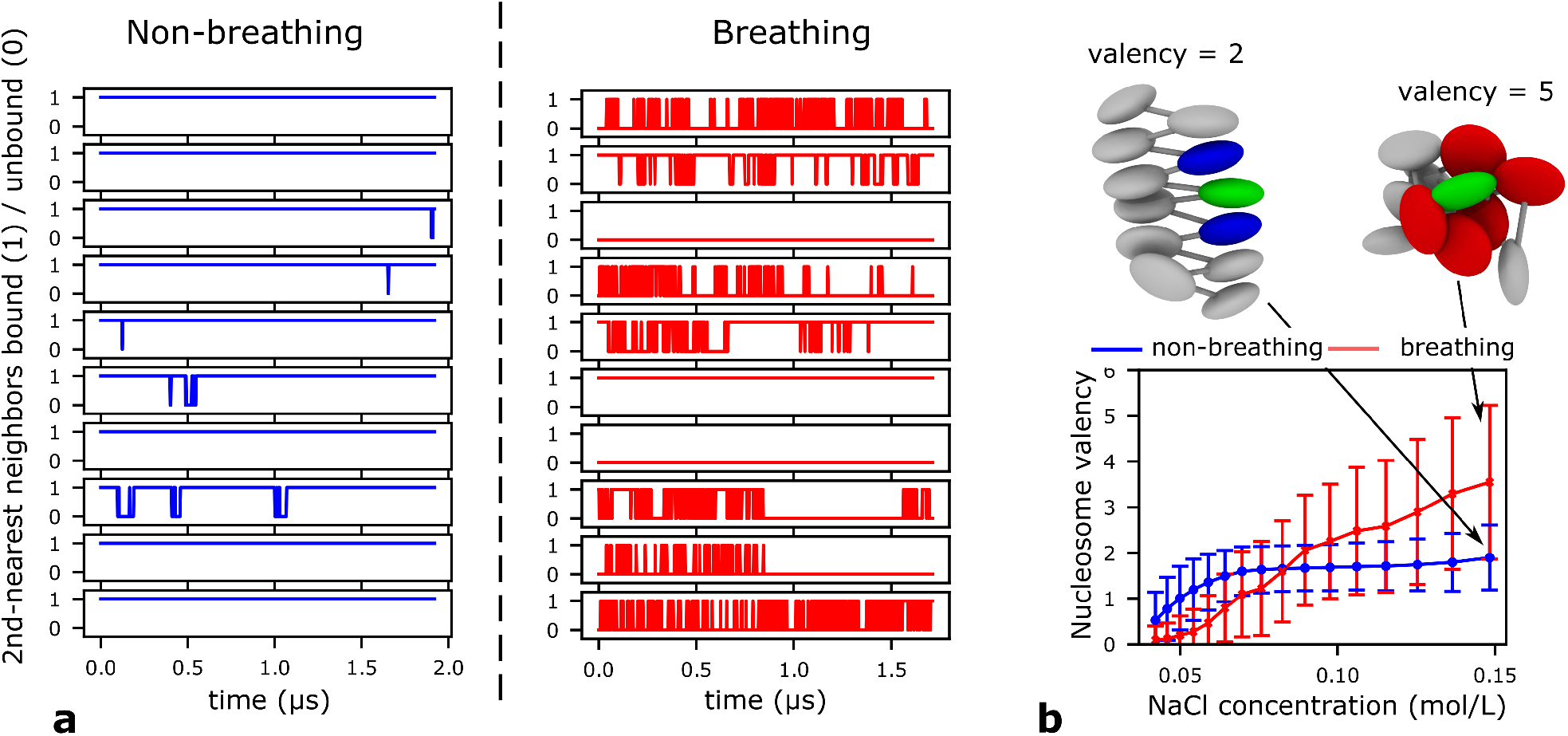
Dynamic nature of nucleosome–nucleosome interactions within compact chromatin. **a** Time series of the binding (1) and unbinding (0) of second-nearest nucleosome neighbor pairs computed from ten independent unbiased MD trajectories at 0.10 mol/L NaCl, and comparing the behavior of non-breathing (left; blue) versus breathing (right; red) nucleosomes. **b** Average valency of a non-breathing (blue) versus a breathing (red) nucleosome—defined as the number of nucleosome neighbors that are ‘in contact’ (i.e., within a center-to-center distance of 110 Å) of itself (see Supporting Information)—within compact chromatin versus NaCl concentration. The cartoon at the top-left highlights a non-breathing reference nucleosome (green) with a valency of two (i.e., in contact with the two blue nucleosomes). The cartoon at the top-right illustrates a breathing nucleosome (green) that is in contact with five other nucleosome neighbors (red).

A fascinating insight stemming from our simulations is that the molecular interactions sustaining the compact state of liquid-like chromatin are instead highly heterogeneous (Figure 4c,d) and transient (i.e., nucleosomes bind and unbind dynamically; Figure 5b); being most strongly contributed by non-specific electrostatic interactions between the various disordered histone tails and the DNA (both nucleosomal and linker DNA). Hence, unlike in the 30-nm fiber, the important acid patch region within liquid-like chromatin is free to recruit the wide-range of chromatin binding factors that are present in cells^112^. Such diverse short-lived interactions are consistent with chromatin dynamically transitioning between highly heterogeneous compact structures that make up the liquid-like ensemble. Furthermore, the key importance of histone tail–DNA electrostatic interactions is supported by experiments demonstrating that chromatin with tail-less nucleosomes fails to condense^113^. In addition, mutation of all H4 arginine and lysine residues to alanine, or histone tail acetylation inhibits nucleosome–nucleosome interactions and the intrinsic LLPS of chromatin^36^.

Interestingly, despite the huge differences in the molecular interactions that stabilize the liquid-like and the fiber-like structures of chromatin, the dominant role of H4K16 is uncontested in both cases (Figure 4). Among the contacts established by the H4 tail, the electrostatic interactions mediated by K16 are the strongest (most significantly with DNA in both types of chromatin structures, and more modestly with the acidic patch in the zigzag fibers). This dominance of the H4K16 residue is in agreement with the well-known decompaction triggered by H4K16 acetylation^113,114^, and the observation that reversible acetylation of H4K16—one of the most frequent post-translational modifications across organisms—has diverse functional implications^115^.

Besides characterizing the molecular features of liquid-like chromatin, we were interested in understanding the underlying physical principles that drive chromatin to adopt such a disordered organization, in terms of its thermodynamics. The structural heterogeneity of the liquid-like ensemble (Figure 3c) indicates that such organization decreases the free energy of chromatin by expanding the number of accessible microstates (entropy gain), compared to those available in a regular 30-nm fiber. Next, to assess the variation in enthalpy, we define the ‘valency’ of nucleosomes as the average number of nucleosome–nucleosome contacts (see Supporting Information). We observe that nucleosomes within liquid-like chromatin have a significantly higher valency at physiological conditions than nucleosomes within 30-nm fibers (Figure 5b). Hence, beyond the entropic driving force, nucleosomes within a liquid-like ensemble decrease their free energy by establishing numerous, but still transient, attractive interactions that maximize the enthalpy gain upon chromatin compaction. Numerous weak and transient internucleosome contacts are preferred over fewer strong longer-lived face-to-face interactions, as the latter represents a greater entropic cost.

### Physical and molecular determinants of intrinsic liquid–liquid phase separation of chromatin

Recent groundbreaking experiments discovered that 12-nucleosome reconstituted chromatin undergoes intrinsic LLPS—i.e., without the aid of additional proteins—under physiological salt concentrations in vitro and when microinjected into cells^36^. Extensive studies characterizing the LLPS of proteins and nucleic acids have demonstrated that multivalency is the dominant driving force for their LLPS^50–52,116–118^. That is, proteins with high valencies can stabilize LLPS by forming numerous weak attractive protein–protein^50,51,116^, protein–RNA^119^, and/or protein–DNA^33,34^ interactions that compensate for the entropy loss upon demixing^52^. Furthermore, binding of multiple Swi6 (the Schizosaccharomyces pombe HP1 protein) molecules to nucleosomes was recently shown to reshape the nucleosome in a manner consistent with nucleosome breathing—i.e., increasing the solvent exposure of core histones—and subsequently promote HP1-chromatin LLPS^45^. These ideas, together with our observations of nucleosome valency enhancement from spontaneous breathing, led us to hypothesize that such nucleosome plasticity is crucial in facilitating the intrinsic LLPS of chromatin.

To investigate this phenomenon and gain new molecular and thermodynamic insight, we use our minimal coarse-grained chromatin model that can simulate a solution of hundreds of interacting chromatin arrays. Specifically, we perform direct coexistence simulations of systems containing 125 independent 12-nucleosome chromatin arrays with a uniform NRL of 165 bp at different conditions. The direct coexistence method involves simulating two different phases—the condensed (chromatin-enriched) liquid in contact with the diluted (chromatin-depleted) liquid—in the same simulation box separated by an interface^120–122^ (see Supporting Information). From these simulations, we compute full liquid–liquid phase diagrams of chromatin at constant room temperature in the ‘NaCl concentration’ versus ‘chromatin density’ space (see Supporting Information and Figure S5), and compare the results for chromatin with breathing nucleosomes and, as a control, for nucleosomes that are artificially constrained to remain fully wrapped (i.e., non-breathing).

Our phase diagrams allow us to compare the conditions under which chromatin LLPS takes place spontaneously for the two representations. This analysis reveals that, besides sustaining the liquid-like behavior of individual chromatin arrays, one immediate effect of the enhancement of nucleosome valency by nucleosome thermal fluctuations is increasing the range of stability of the intrinsic LLPS of chromatin (Figure 6a). That is, chromatin spontaneously forms condensates above a critical monovalent salt concentration; i.e., where screening by counterions is sufficiently strong to eliminate the DNA–DNA repulsion and encourage the formation of numerous weak and transient attractive internucleosome interactions. Compared with breathing nucleosomes, when nucleosomes are constrained to remain fully wrapped, chromatin phase separation requires higher NaCl concentrations to become thermodynamically stable (Figure 6a); the limited valency of non-breathing nucleosomes necessitates the formation of stronger nucleosome–nucleosome interactions to obtain a sufficient enthalpic gain for LLPS. Beyond the critical solution salt concentration, the size of the liquid–liquid coexistence region is significantly larger for chromatin with nucleosomes that breathe spontaneously (Figure 6a), which implies that under the same solution conditions, breathing nucleosomes yield a more dense and, hence, more stable condensed liquid.

**Figure 6.**
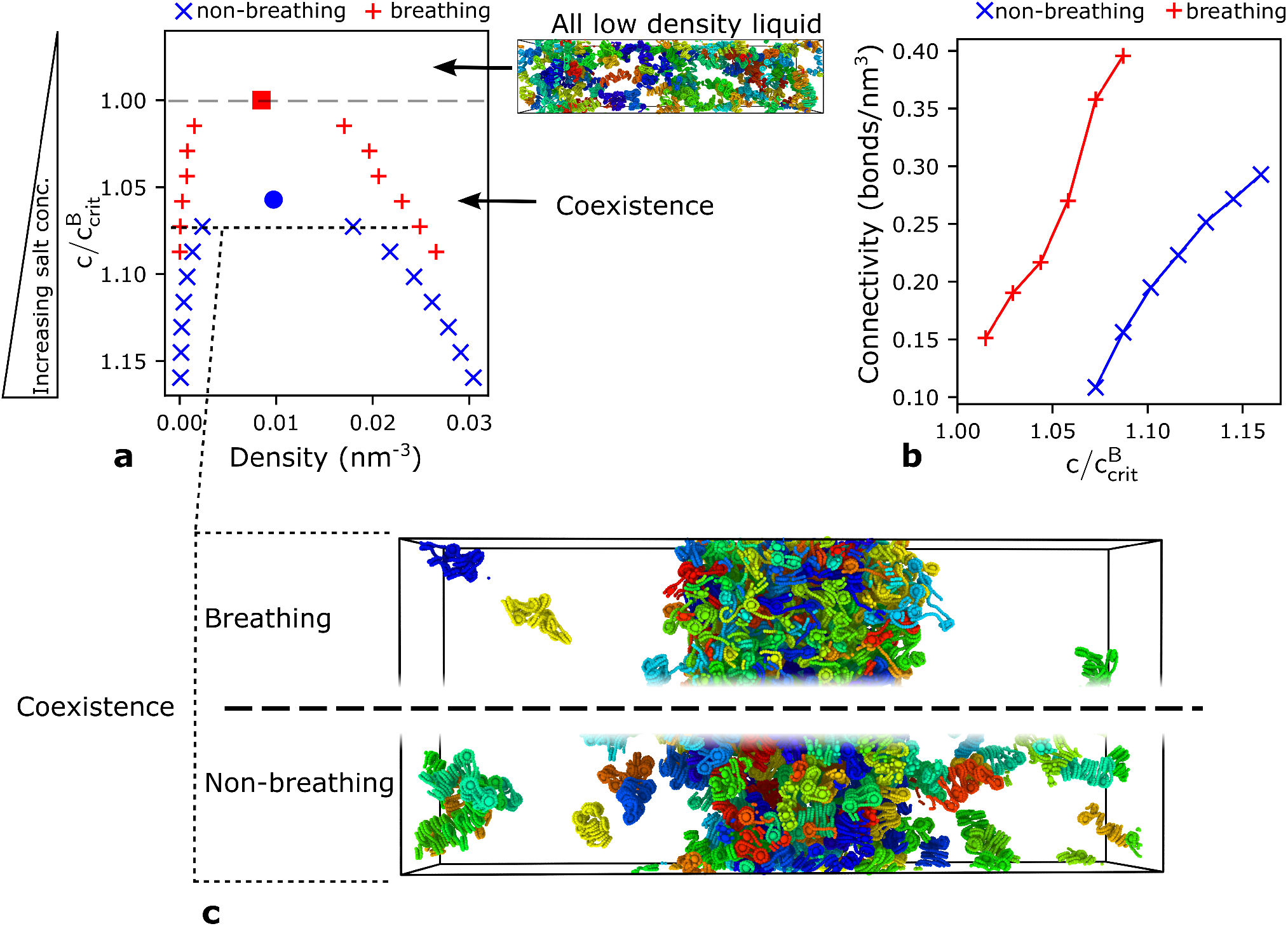
Impact of nucleosome breathing in the phase behavior of chromatin. **a** Phase diagram of a solution of 12-nucleosome chromatin arrays exploring the space of decreasing concentration of NaCl (vertical axis, *C*) versus density (horizontal axis). The datapoints (blue: non-breathing, red: breathing) represent coexitence points, i.e., the densities of the chromatin-diluted (left branch) and chromatin-enriched (right branch) coexisting liquid phases at a given monovalent salt concentration. The density is defined as the number of chromatin molecules per unit of volume in *nm*^−3^. The region above the coexistence curve is the ‘one-phase region’, where the NaCl concentration is too low to screen DNA–DNA electrostatic repulsion and enable the attractive nucleosome–nucleosome interactions, needed for LLPS (see top snapshot of a well-mixed low density liquid). The region below the coexistence curve represents the ‘two-phase region’ where chromatin demixes into a condensed (chromatin-enriched) and a diluted (chromatin-depleted) liquid phase. The maximum in the coexistence curve represents the critical point: if the NaCl concentrations exceeds the critical value, LLPS occurs spontaneously. The vertical axis has been normalized by the critical salt concentration for chromatin with breathing nucleosomes 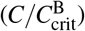, where 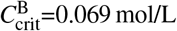 mol/L of NaCl. The critical salt concentration for chromatin with non-breathing nucleosomes is 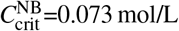 mol/L. **b** Connectivity of the condensed liquid formed by chromatin with non-breathing (blue) versus breathing (red) nucleosomes. The connectivity is defined as the mean number of connections that chromatin arrays within the condensed liquid form (i.e., the number of distinct chromatin arrays that a reference array is in contact with, considering any two nucleosomes within a center-to-center distance shorter than 110 Å) multiplied by the density of the condensed phase. **c** Representative simulation snapshots of the phase-separated liquids formed by chromatin with breathing (top) versus non-breathing (bottom) nucleosomes at a value of 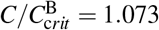 (i.e., *C*=0.074 mol/L of NaCl).

To analyze the physical forces governing LLPS of chromatin, we compute the average density of connections (i.e., bonds) that nucleosomes form per unit of volume within the condensed phase at a fixed salt concentration. Similar to its effect in enhancing the valency of nucleosomes within single chromatin arrays, we observe that nucleosome breathing promotes LLPS because it fosters a much higher nucleosome–nucleosome connectivity within the condensed liquid (Figure 6b). A densely connected chromatin liquid phase fulfills two crucial requirements: making condensation thermodynamically favorable and maintaining the liquidity of the condensed phase. A condensed chromatin liquid and a gel would exhibit the same type of percolating network structure (i.e., where each chromatin array is bound transiently to at least one other), with increasingly strong and long-lived nucleosome–nucleosome interactions favoring a dynamically arrested gel-like state. In other words, because nucleosome–nucleosome interactions must remain sufficiently weak, and thus short-lived, to preserve the liquid-like properties of the condensed phase, a densely connected network ensures that, collectively, the low strength of nucleosome– nucleosome interactions compensates for the entropy loss upon demixing. Hence, liquid chromatin condensates are favored by the dynamic formation and rupture of a large number of weak attractive nucleosome–nucleosome interactions, which in turn are facilitated by the dynamic breathing behavior of nucleosomes.

## Discussion

We introduce a novel mechanistic multiscale model of chromatin designed to investigate the connection between the fine-atomistic details of nucleosomes and the emergence of chromatin self-organization and LLPS in systems with over a thousand nucleosomes. By pairing a residue/base-pair resolution model with a minimal representation of chromatin, our multiscale approach enables the study of collective effects of amino acid mutations, post-translational modifications, histone secondary structural changes, DNA sequence, and nucleosome dynamics in modulating mesoscale chromatin structural properties.

Our simulations put forward nucleosome plasticity at physiological salt conditions (which can be driven not only by nucleosome breathing, but also by sliding, HP1 binding, nucleosome remodeling, post-translational histone modifications, histone replacement, and other mechanisms) as a key driving force of the intrinsic liquid-like behavior of chromatin. Nucleosome plasticity transforms nucleosomes from the uniform and static disc-like repeating units needed to sustain rigid fibers, to highly heterogeneous and dynamical particles that engage in promiscuous nucleosome–nucleosome interactions, sample a wide range of internucleosome rotational angles, and spontaneously self-assemble into disordered liquid-like structures. Unlike the 30-nm fibers which are strongly contributed by H4 tail to acidic patch interactions, liquid-like chromatin is instead stabilized by a diverse range of non-specific DNA–histone tail electrostatic interactions. Therefore, within liquid-like chromatin the nucleosome acidic patch region can be more easily accessed by the myriad of chromatin-binding factors that have been proposed to target it in vivo^112,123^, which include HP1^124^. Furthermore, controlled access to the acidic patch region has been hypothesized to play a crucial role in the modulation of chromatin remodelling motors that regulate nucleosome sliding^125^.

Importantly, our work demonstrates that liquid-like chromatin is simultaneously stochastically organized and tightly packed. This suggests that chromatin compaction does not immediately imply steric barring of enzymes, and that indeed chromatin as a compact liquid-like system is optimum for DNA searchability; i.e., liquid-like chromatin provides easier and more homogeneous DNA access for processes like ‘scanning and targeting genomic DNA’ without the need for chromatin to undergo decompaction, as postulated by Maeshima and collaborators^18^. Therefore, the strong link between the stochastic organization of chromatin— which we show might facilitate access of the transcriptional machinery to nucleosomal DNA—and nucleosome plasticity might have important functional implications. Nucleosomes represent a fluctuating barrier for the binding of transcription factors to DNA, and hence, for transcription^66,67^; the maximally repressive state is that of the fully wrapped nucleosome, while the maximally non-repressive state is that of a nucleosome-free region^126^. However, although the majority of transcription factors seem to bind to nucleosome-free DNA regions^127,128^, spontaneous nucleosome breathing has been suggested to provide transient access to some of these proteins to their nucleosome binding sites and, hence, facilitate transcription initiation^129–131^, specially at physiological ionic strengths^60^. The increased access of chromatin due to nucleosome breathing might be most relevant for rationalizing the mechanisms of binding of pioneering factors^132^, which are a special class of proteins that can secure their target DNA sites on nucleosomes and in compact chromatin^133,134^. Additionally, in vitro, nucleosome breathing has been suggested to play a role in transcription elongation, by facilitating the movement of RNA polymerase II ternary elongation complex across the nucleosomal DNA^66,135^. In vivo, spontaneous nucleosome breathing, combined with the action of chromatin remodellers, allows the Clustered Regularly Interspaced Short Palindromic Repeats (CRISPR) associated protein 9 (Cas9) to access the nucleosomal DNA^136^. Our observations also support one potential mechanism influencing control of gene transcription kinetics: spontaneous thermal breathing. This breathing behavior has been proposed to allow nucleosomes to adopt a wide-range of promoter configurations, some of which transiently facilitate transcription and others that momentarily inhibit it; hence, modulating the rate of discontinuous transcription of genes that includes bursts of activity^126^. That is, the required malleability of promoter configurations is consistent with the high flexibility, structural heterogeneity, and potential for LLPS of chromatin with dynamical nucleosomes stemming from our work.

The attractive interactions that maintain the DNA wrapped around the histone core are predominantly electrostatic in nature; hence, the breathing motion of nucleosomes is boosted by electrostatic screening at physiological salt concentrations. Experiments have demonstrated that the low to moderate salt concentrations used in many in vitro experiments (up to ∼0.08 mol/L of NaCl), hinder nucleosome breathing (≤10%), while physiological salt conditions (above ∼0.1 mol/L of NaCl) promote it^59^. Our work further reveals that significant nucleosome breathing at physiological salt favors the liquid-like behavior of chromatin even in artificially homogeneous oligonucleosomes (e.g., with uniform DNA linker lengths and DNA sequences and regular histone compositions). In contrast, inhibiting nucleosome fluctuations drives chromatin to form ordered 30-nm zigzag fibers. Accordingly, our work suggests that the modulation of nucleosome breathing with salt can aid in reconciling discrepancies between the ordered and disordered chromatin structural models derived from in vitro and in vivo experiments, respectively.

Significantly, we demonstrate that the same molecular and biophysical driving forces that sustain the liquid-like behavior of nucleosomes within single compact chromatin arrays also promote the intrinsic phase separation of a solution of chromatin arrays. Specifically, nucleosome breathing fosters not only the formation of disordered and flexible chromatin ensembles, but also LLPS and the emergence of phase-separated chromatin compartments. Interestingly, when nucleosome breathing is artificially suppressed (analogous to the behavior of strongly positioning artificial sequences), chromatin shows reduced propensity for LLPS. Thus, intranuclear conditions that can spontaneously tune intrinsic nucleosome breathing via modulation of electrostatic interactions (e.g., changes in ionic salt concentration, pH, DNA/histone mutations) are expected to alter the spatial organization, degree of compaction, and compartmentalization of chromatin. Modulation of nucleosome breathing may, therefore, represent a key mechanism used by cells, not only to organize chromatin but also, to modulate its function.

Our work also reveals that nucleosome breathing and multivalency are intricately linked and are positively correlated. Multivalency has been previously identified as an important property governing protein LLPS, both in the cytoplasm and nucleoplasm. Furthermore, protein multivalency and, therefore, LLPS can be modulated by several external (e.g., salt concentration, temperature, pH, multi-component composition) and intrinsic factors (e.g., protein mutations, post-translational modifications, and disorder-to-order transitions). For chromatin, we show that increase in monovalent salt concentration facilitates nucleosome breathing that directly enhances nucleosome multivalency. Moreover, our work strongly suggests that, in the absence of significant environmental changes, within a fiber-like chromatin model, the valency of nucleosomes can be considered as roughly static; i.e., the torsional rigidity of the DNA locks nucleosomes in a face-to-face arrangement, where they bind to one another strongly. In contrast, within a plastic nucleosomal framework, nucleosomes, like proteins, possess an inherent capacity (i.e., nucleosomal breathing) to dynamically modify their valency, and ultimately regulate their functionality. Recent landmark experiments by Sanulli et al.^45^ reveal that the binding of many HP1 molecules unexpectedly reshapes nucleosomes and makes the histone core more accessible to the solvent, consistent with nucleosomal DNA unwrapping, which in turn can increase the availability of the core for interactions that facilitate LLPS. Our work further suggests that the increase in the plasticity of nucleosomes induced by HP1, is consistent with the amplification of: (a) the local flexibility of chromatin, (b) the range of accessible nucleosome–nucleosome pair orientations, (c) the effective valency of nucleosomes, and (d) the transient nature of the inter-nucleosomal attractive interactions that lead to chromatin LLPS.

Together, our work postulates that nucleosome plasticity is an intrinsic property of nucleosomes that facilitates chromatin stochastic self-assembly and LLPS, and hence, contributes to regulate the organization and membraneless compartmentalization of the genome. These findings advance the molecular mechanisms and biophysical understanding of how the liquid-like organization of the genome is formed and sustained, and how it can be regulated. Modulation of nucleosome plasticity might have important implications in the functional organization of the genome and in the control of gene transcription parameters.

## Methods

### Chemically-specific chromatin coarse-grained model

Our chemically-specific chromatin model represents every residue—both within the histone globular regions and histone tail regions—explicitly using a single bead that carries the charge, hydrophobicity, and size of its atomistic counterpart, following the work of Dignon et al.^91^. Each bead is defined as a point particle with an excluded volume centered on the *C*_*α*_ atom of the amino acid it represents. For each amino acid, the bead diameter is calculated from the experimentally-measured van der Waals volumes and assuming that amino acids have a spherical shape, as done previously^89^. We also assign a charge to each bead corresponding to the total charge of the related amino acid. The sequence-dependent hydrophobic attraction between specific amino acid pairs is accounted for by the Kim–Hummer model^89,91^, which consists of a shifted and truncated Lennard-Jones potential with parameters derived from experimental amino-acid pairwise contact propensities^90^. Using our previous microsecond-long Bias-exchange metadynamics molecular dynamics simulations of 211-bp nucleosomes^31^, we differentiate between the globular histone core, residues that retain their secondary structure throughout the simulation, and the disordered histone tails. We then treat amino acids within intrinsically disordered regions as fully flexible polymers (i.e., with no energetic penalty for bending) using a harmonic potential with a stiffness bond constant *k*_*b*_ of 10 kcal/mol/Å^2^ and a resting length *r*_0_ of 3.5 Å, as proposed by Dignon et al.^91^. Because the amino acids within the globular histone core regions, which are largely *α*-helical, exhibit relatively small structural fluctuations when compared to those of histone tails in our simulations^31^, we describe their structural fluctuations by building an elastic network model^137^, which avoids the need of including internal non-bonded terms in these regions. In practice, we take a representative structure from the highest populated cluster in our atomistic simulations^31^ as the reference structure, and connect all the globular histone core beads that are within 7.5 Å of each other with harmonic springs and form a Gaussian elastic network model (GNM)^137^. For the harmonic bond interaction of the GNM, we use a spring constant *k*_*GNM*_ of 10 kcal/mol/Å^2^, and the equilibrium distance among amino-acid pairs is set equal to its value in the reference atomistic structure.

The second crucial feature of our model is that it considers sequence-dependent DNA mechanical properties by using the Rigid Base Pair (RBP)^84–88^ model with added phosphate charges. The RBP model represents DNA base-pair steps with one ellipsoid, and models DNA conformational changes in terms of harmonic deformations of six helical parameters (three angles: twist, roll, and tilt, and three distances: slide, shift, and rise) that account for the relative orientations and positions of neighboring base-pair planes. The DNA mechanical potential energy is computed from the sum of harmonic distortions of equilibrium base-pair step geometries. We used the Orozco group parameters^87,88^ computed from MD atomistic simulations— i.e., the equilibrium values by fitting Gaussian functions to the distributions of helical parameters, and the elastic force constants by inversion of the covariance matrix in helical space. In practice, our model represents single base pairs, both within the nucleosomal and linker DNA, by one coarse-grained bead defined by a position vector, ***r***, and an orientation quaternion, *q*. We add two virtual charge sites to each DNA ellipsoid (i.e., one per phosphate approximating the shape of the DNA phosphate backbone) to consider the crucial electrostatic interactions that drive chromatin self-organization. We implement this RBP plus charged virtual sites in LAMMPS (http://lammps.sandia.gov)^138^ with an ellipsoid defined by two-point particles with a negligible but non zero mass. While the combined three-particle base-pair bead is treated as a single rigid body for the dynamics, the individual components each contribute to the calculation of the potential and forces.

Besides the excluded volume and hydrophobic attraction, we consider electrostatic interactions between the charged beads by means of a Debye–Hückel potential that approximates screening by monovalent counterions in solution. Although this potential makes several approximations, like not accounting for ion condensation and ion-ion correlation effects, it has been shown to capture well the salt-dependent compaction of chromatin at the relatively low ionic concentrations present inside cells (≤ 0.15 mol/L of NaCl)^16,20^. While the Debye-Hückel approximation is exact in the low salt limit, its numerical implementation becomes progressively challenging at low salt because estimating interactions with higher Debye-lengths requires using larger cutoffs (see Supporting Information). As the final ingredient, we omit non-bonded interactions between immediately bonded beads.

Binding of the nucleosomal DNA to the histone protein core is achieved through these protein–DNA electrostatic interactions, resulting in a nucleosomal DNA that wraps ∼1.7 times around the histone core and exhibits spontaneous unwrapping/re-wrapping. As discussed in the ‘Model Validation’ and Supporting Information, the force-induced unwrapping behavior of these nucleosomes is in quantitative agreement with experiments at single base-pair resolution.

To model nucleosomes that are artificially constrained to be ‘non-breathing’, we describe the histone core together with the bound nucleosomal DNA as a single GNM, using the same 7.5 Å threshold and bond parameters. This results in the nucleosomal DNA being constrained to remain permanently bound to the histone core, and inhibits both nucleosome breathing and sliding.

In summary, our chemically-specific coarse-grained model of chromatin preserves the shape and size of the nucleosome core, the differential fluctuations between the globular and histone tail regions, the length of the histone tails and their flexibility, and the explicit charges in the histone tails and on the nucleosome surface, including the acid patch region^3^, the differential hydrophobic attraction among distinct amino acids, the sequence-dependent mechanical properties of the DNA, and the thermal breathing of nucleosomes.

### Minimal chromatin coarse-grained model

When devising our minimal model, we took care in capturing the manner in which nucleosomes interact with one another and the mechanical properties of the DNA; including, its torsional and bending flexibility that we observe in our chemically-specific chromatin simulations. Both features are crucial in defining the conformational space of nucleosomes; indeed, if we ignore the DNA torsion and model DNA simply as a worm-like chain we observe a spurious nucleosome organization (not shown). To account for these two features, while enabling efficient sampling of a few thousand nucleosomes, our minimal chromatin model approximates the shape and size of the histone core with a single bead (using an ellipsoid of 28 *×* 28 *×* 20 Å radii), and represents linker and nucleosomal DNA with one finite-size orientable sphere (using an ellipsoid of 12 *×* 12 *×* 12 Å radii) for every 5 base-pair steps. Inspired by the success of the RBP model in adequately approximating the atomistic mechanical properties of DNA at a single base-pair-step resolution, we propose a ‘minimal RBP-like model’ for DNA at a resolution of 5-base-pairs per bead. Accordingly, we optimize the minimal helical parameters from chemically-specific coarse-grained simulations of 200 bp DNA strands (see Supporting Information). Then we describe nucleosome–nucleosome and nucleosome–DNA interactions by a series of orientationally-dependent potentials fitted to reproduce internucleosome potentials of mean force calculated with our chemically-specific coarse-grained chromatin model (see Figure S4 and Supporting Information). To represent breathing nucleosomes, we analyze our chemically-specific chromatin simulations to determine the fraction of DNA that remains predominantly bound to the histone core, and define only that fraction as nucleosomal DNA. For each snapshot in the chemically-specific model trajectories we will get a different definition which builds up a set of structures representative of the thermodynamic equilibrium distribution of nucleosome breathing states. By using multiple different structures from this set we can incorporate the effects of nucleosome breathing in the model without having to directly simulate completely free DNA, significantly reducing the degrees of freedom, enabling superior sampling. In the case of the non-breathing nucleosomes we simply define the nucleosomal DNA following the same definition used in the chemically-specific model. Then, in both cases, breathing and non-breathing nucleosomes, we attach permanently the nucleosomal DNA to the histone core bead. Hence breathing and non-breathing nucleosomes are constructed using equilibrium configurations from different chemical-specific simulations (Figure S3). In addition, to account for the slightly exposed histone core in breathing nucleosomes, we add an additional anisotropic potential to the total energy. We parameterize this anisotropic term to be consistent with experimental force spectroscopy experiments on single nucleosomes^100–103^ (see Supporting Information, and Figures S2 and S4).

### Debye-length Hamiltonian replica exchange

To drive chromatin systems over the free-energy barriers and achieve complete sampling, we developed a Hamiltonian exchange method that attempts exchanges between replicas with different values of the Debye-length of the screened-coulomb interaction (*λ*_*D*_) while keeping the temperatures the same. As in standard Hamiltonian exchange, the exchange probability is given by:

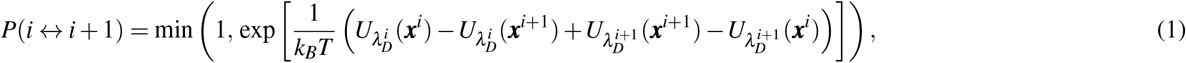

where ***x***^*i*^ are the chromatin coordinates of the *i*^th^ replica, and 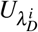 the potential energy function at Debye length 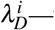—the original Debye length of the *i*^th^ replica. The exchange is accepted or rejected based on the Metropolis criteria, and upon exchange the potential energy functions (or coordinates) are switched. For our 12-nucleosome chromatin systems, we find that at the value of *λ*_*D*_ = 8.0 Å (corresponding to 0.15 mol/L salt), the chromatin structures are compact and suffer from sampling issues, increasing the Debye length to 15 Å gives rise to open structures that can sample effectively. We find that a range of 8.0–15.0 Å requires 16 replicas to get an acceptance probability of 30 %. This is significantly less than the ∼80 replicas that we found would be required for standard temperature replica exchange to sample the range of 300 K to 600 K with a similar exchange probability. Additionally the different Debye lengths are all at physically relevant salt concentrations, therefore while increasing the sampling we can investigate the salt dependent behavior of the system. All replicas give us useful information, whereas for temperature replica exchange only the replica at the target temperature can be used for analysis.

## Acknowledgements

This project has received funding from the European Research Council (ERC) under the European Union’s Horizon 2020 research and innovation programme (grant agreement No 803326). R.C.-G. is an Advanced Fellow from the Winton Programme for the Physics of Sustainability. S.E.F. would like to acknowledge the EPSRC Centre for Doctoral Training in Computational Methods for Materials Science for funding under grant number EP/L015552/1. E.J.W. is funded by an EPSRC studentship from grant EP/R513180/1. J.A.J. is a Junior Research Fellow at King’s College. This work has been performed using resources provided by the Cambridge Tier-2 system operated by the University of Cambridge Research Computing Service (http://www.hpc.cam.ac.uk) funded by EPSRC Tier-2 capital grant EP/P020259/1.

## Author contributions statement

R.C.-G. and S.E.F. conceived the project; S.E.F, E.J.W., and R.C.-G. designed the models and the computational framework; S.E.F. performed research and analyzed data; A.G. contributed to the computational implementation of the model; J.A.J. contributed to the interpretation and contextualization of the data; R.C.-G. wrote the manuscript with help from S.E.F, J.A.J., and E.J.W.; All authors reviewed the manuscript; R.C.-G. acquired funding; and R.C.-G. supervised research.

## Additional information

The authors declare no competing interests.

## Supporting Information

### I. CHROMATIN MULTISCALE METHODOLOGY

We develop a multiscale method to investigate self-organization and liquid–liquid phase separation (LLPS) of chromatin at a range of resolutions, and to link atomistic properties with the emergence of collective behavior (Figure 1, S1). In previous work, we performed simulations at level 1—all-atom simulations of a 211-bp nucleosome in explicit solvent with ions—that enabled the differentiation between histone globular domains and the histone tails [1]. Information was distilled from level 1 to develop level 2—our ‘chemically-specific’ coarse-grained model, which considers amino-acid and DNA base-pair level resolution. To reach the system sizes needed to investigate LLPS of chromatin arrays, further coarse-graining was developed to produce level 3—our ‘minimal’ coarse-grained model at nucleosome level resolution.

**Figure S1.**
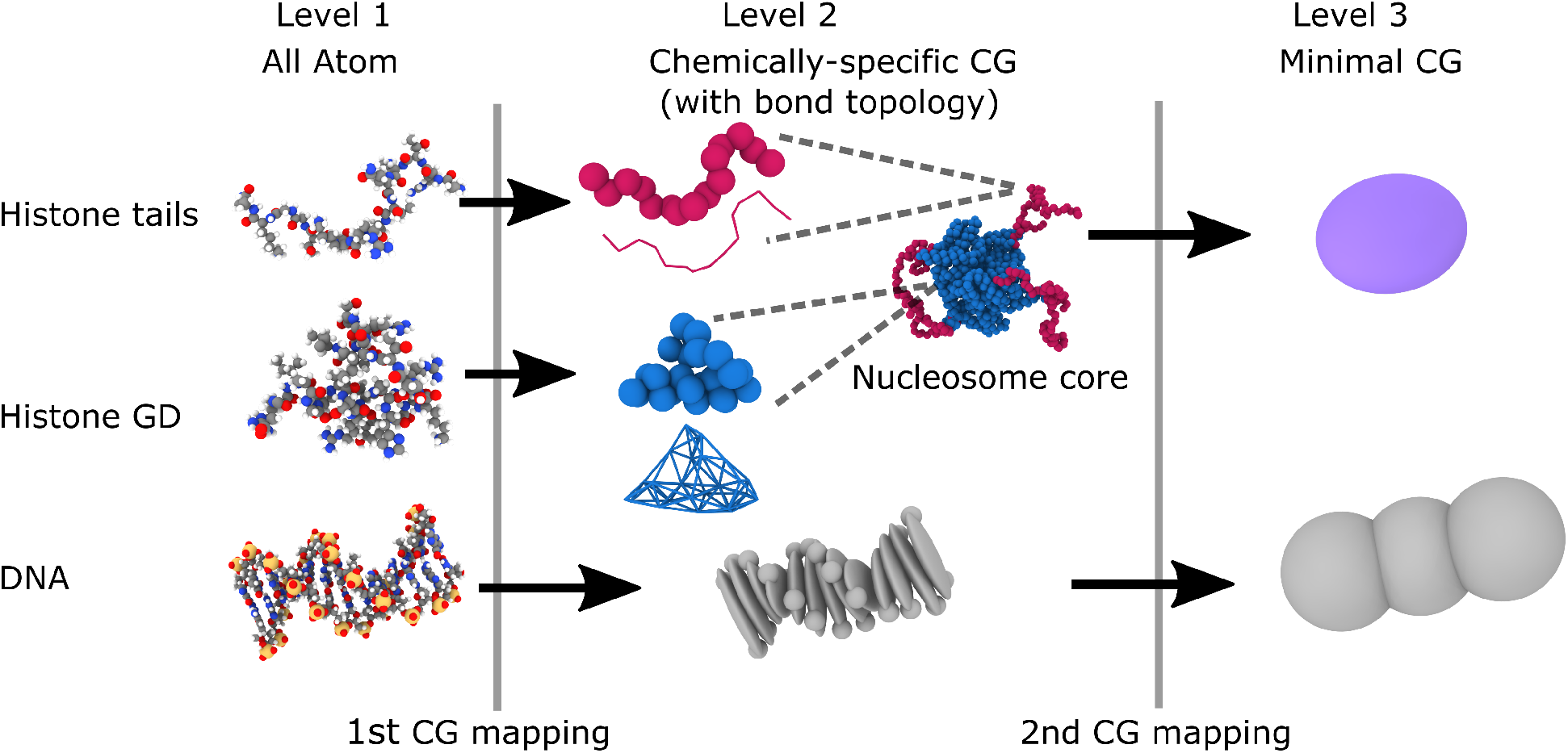
Additional details on the multiscale coarse-grained mapping, complimentary to Figure 1. The three panels provide schematic representations of the different particles featured in our multiscale approach, which spans three levels of resolution. **(Level 1)** All-atom representations of histone tails, histone globular domains (GD), and DNA. **(Level 2)** Chemically-specific representations of histones, both tails and GD, with the difference in bond topology among them illustrated, and DNA. **(Level 3)** Minimal coarse-grained model showing the histone core and DNA mapping.

### II. COMPUTATIONAL IMPLEMENTATION

We implement both the chemically-specific and minimal model in the molecular dynamics (MD) package ‘Large-scale Atomic/Molecular Massively Parallel Simulator’ (LAMMPS; https://lammps.sandia.gov/) [2]. Finite size ellipsoidal particles—essential for the implementation of orientation dependent potentials—from the `ASPHERE` package are used to represent coarse-grained segments of chromatin. Additionally, the `RIGID` package is used to enable the building of composite rigid bodies.

We use LAMMPS stable version 3rd March 2020 compiled with our custom code, which is available upon request. All simulations were performed on the Cambridge Service for Data Driven Discovery (CSD3).

### III. CHEMICALLY-SPECIFIC COARSE-GRAINED MODEL

#### A. Energy function of chemically-specific model

The total potential energy function for the chemically-specific coarse-grained model is,

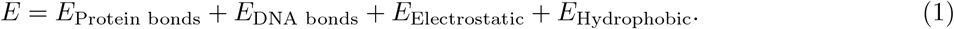

Both the electrostatic and hydrophobic terms have a cutoff distance—beyond which the energy goes to zero— which is necessary for efficient computational implementation. The protein bonds are a standard harmonic interaction,

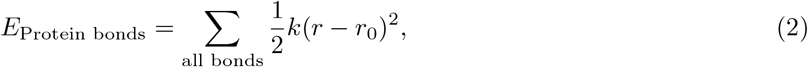

where *k* = 10 kcal*/*mol*/*Å^2^ is the spring constant and *r*_0_ is the (bond dependent) equilibrium bond length.

The DNA bonds are modeled by the Rigid Base Pair (RBP) potential [3–7], which represents DNA base-pairs as rigid planes and the interactions between two adjacent base-pairs in terms of harmonic deformations of six helical parameters (three displacements: shift, slide, rise, and three angles: tilt, roll, twist).

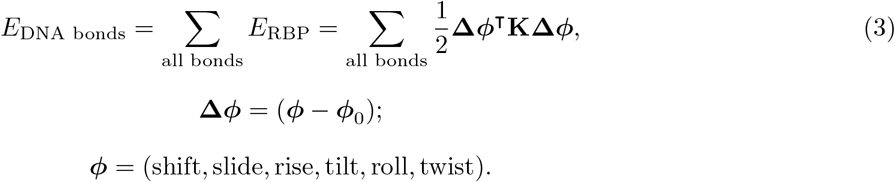

where ***φ*** is a 6-dimensional vector of the 6 DNA helical parameters, ***φ***_0_ are the equilibrium values and **K** is the 6 *×* 6 stiffness matrix. This model is DNA sequence dependent so **K** and ***φ***_0_ are dependent on the base pair step. The helical parameters are determined using the SCHNAaP method [8]. The DNA–DNA bonds use the RBP potential parameters from the Orozco group [6, 7].

Electrostatics are modeled by the Debye-Hückel potential, which accounts for screening due to an implicit monovalent salt solvent,

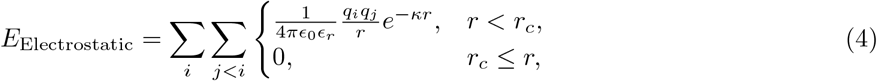

where *υ*_0_ and *υ*_*r*_ are the permittivity of free space and the relative permittivity respectively, *q*_*i*_ and *q*_*j*_ are the charges of particles *i* and *j, r* is the distance between the interacting pair of particles, *κ* is the inverse Debye length and *r*_*c*_ is the cut-off distance (necessary for efficient computational implementation). Electrostatic interactions are considered between all charged particles within the cutoff distance.

To describe short-range non-bonded interactions, we follow the work of Dignon et al. [9] and use their version of the Miyazawa-Jerningan potential [10] with the functional form and parameters of Kim and Hummer [11]. This consists of a sequence-dependent hydrophobic attraction between specific amino acid pairs (Kim-Hummer model [11]), and a standard Lennard-Jones interaction [9].

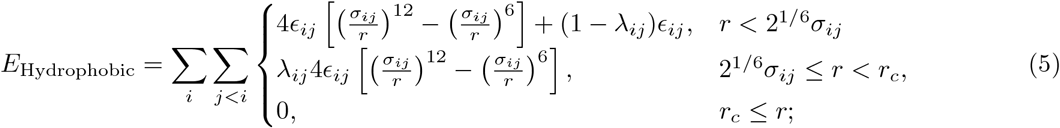

where *υ*_*ij*_ is the dispersion energy, *σ*_*ij*_ is the distance at which a standard Lennard-Jones interaction is zero, *λ*_*ij*_ is a hydrophobic parameter, *r*_*c*_ is the cutoff distance (numerical value different to that for the electrostatic potential) and the sum is over all pairs of particles. Note that when *λ* = 1, we recover a standard Lennard-Jones potential, and when *λ* = 0, we obtain a purely repulsive Weeks-Chandler-Anderson (WCA) potential. Each amino-acid pair has unique values for _*ij*_ and *λ*_*ij*_. *σ*_*ij*_ is defined using the combination rule *σ*_*ij*_ = (*σ*_*i*_ + *σ*_*j*_)*/*2, where *σ*_*i*_, *σ*_*j*_ are the van der Waals radii of particles *i,j* respectively. For interactions involving DNA, *λ*_*ij*_ = 1.

The two non-bonded interactions *E*_Electrostatic_ and *E*_Hydrophobic_ are turned off between directly bonded beads.

#### B. DNA mechanical potential: rigid-base pair forces and torques

Typically the RBP model is used in Monte Carlo simulations, which only require the potential energy and not its derivatives. To implement the RBP potential in MD simulations, forces and torques must be defined. Following the method of Fathizadeh et al [12], the force in the *k*^th^ direction on a base-pair, due to the RBP potential *E*_RBP_ is,

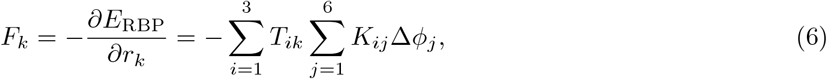

where **T** (see Eq. 45 in Section VII B) is the mid step orientation matrix between the two base-pairs that comprise the bond. **T** can be thought of as the average orientation of the two base-pairs. The torque on a base-pair around the *k*^th^ axis is computed numerically using the central finite difference,

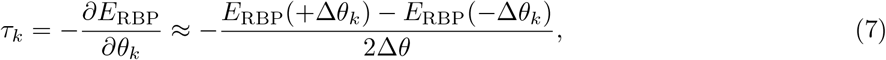

where ∂*θ*_*k*_ represents an infinitesimal rotation about the *k*^*th*^ axis and +Δ*θ*_*k*_ represents rotating the current base-pair about its *k*^*th*^ axis by the small rotation Δ*θ* = 0.00001 rad.

#### C. Model building & Implementation of the chemically-specific model

The chemically-specific coarse-grained model resolves DNA at the base-pair level and proteins at the amino acid level. The DNA beads are modeled using an ellipsoidal particle that approximates the shape and mass of a base-pair. The ellipsoid is rigidly connected to two point particles each with charge -1, which approximates the phosphate backbone and major/minor grooves. The DNA ellipsoids are spatially defined by a position vector ***r***, and a unit quaternion *<u>q</u>* which encodes the orientation and can be concerted into an orthogonal rotation matrix **A**, whose columns are the unit axis vectors of the ellipsoids frame of reference.

We begin by taking our reference nucleosome structure as the most populated structure from our previous all-atom bias-exchange molecular dynamics metadynamics simulations of a 211-bp nucleosome [1]. From there, we define the position and orientation of the DNA ellipsoids by using the software 3DNA [13], which determines the coordinates of the rigid base pairs that fit the atomistic structure. The two phosphate point particles act as virtual charge sites, and for computational reasons have a negligible but non-zero mass of 1 *×* 10^−6^ g/mol. The three RBP particles (the ellipsoid plus the two point charges) are combined together as a rigid body using `fix rigid/nve/small` from the `RIGID` package in LAMMPS. The individual particles within the rigid body each contribute to the resultant potential and forces acting on the combined body, which is treated as a single rigid body for the dynamics.

Using the same atomistic reference structure from our 211-bp nucleosome simulations, the protein beads are defined as point particles centered at the *C*_*α*_ of each atomistic amino acid. The identity of the amino acid type is preserved by defining a different particle type for each unique amino acid. Each amino acid was classified as either belonging to a globular domain or histone tail (Table I). The sequence-dependent mass, charge, van de Waals radius and hydrophobicity values for each amino acid type are given in Table II, which are taken from Ref. [9]. The classification was performed by estimating the persistence of protein secondary structures from our 211-bp nucleosome simulations [1].

**TABLE I.** Definition of histone tail regions. All other residues are classified as belonging to the central globular region.

| Histone | Tail region residues |
| --- | --- |
| H3 | 1-40 |
| H4 | 1-25 |
| H2A | 1-20, 114-128 |
| H2B | 1-25 |

**TABLE II.**
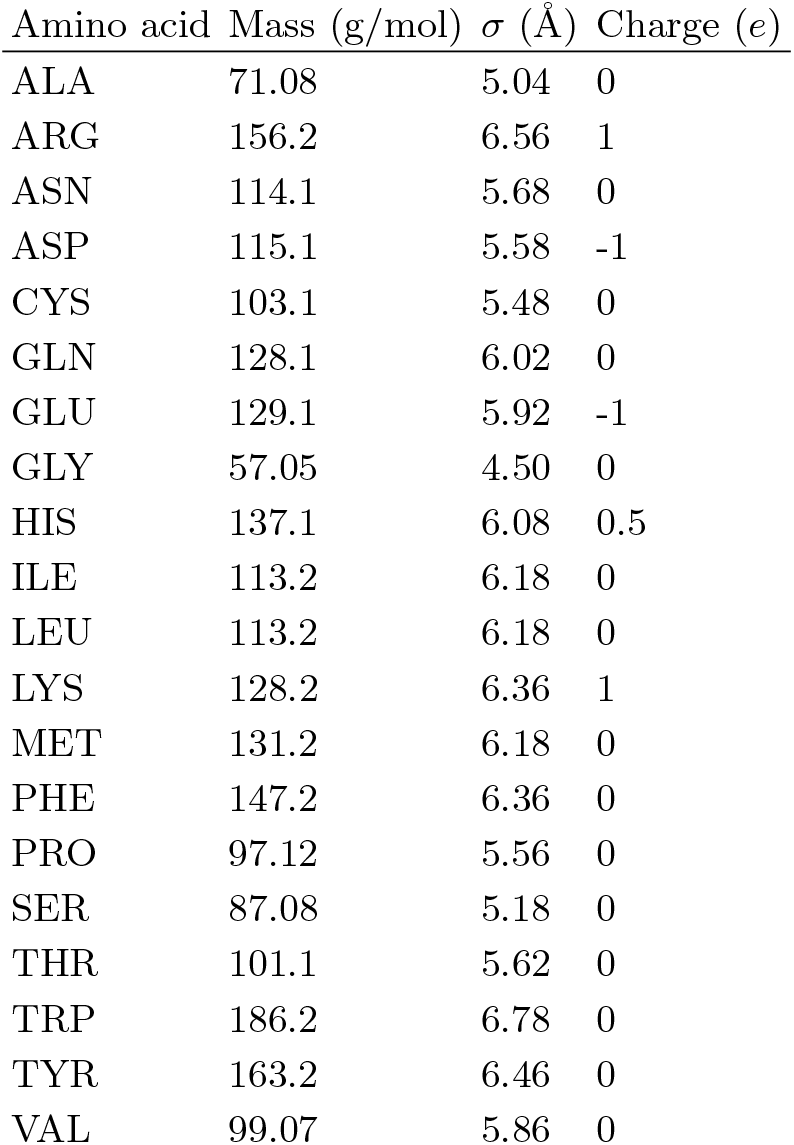
Amino acid parameters taken from Ref. [9].

| Amino acid | Mass (g/mol) | $\sigma$ (Å) | Charge ($e$) |
| --- | --- | --- | --- |
| ALA | 71.08 | 5.04 | 0 |
| ARG | 156.2 | 6.56 | 1 |
| ASN | 114.1 | 5.68 | 0 |
| ASP | 115.1 | 5.58 | -1 |
| CYS | 103.1 | 5.48 | 0 |
| GLN | 128.1 | 6.02 | 0 |
| GLU | 129.1 | 5.92 | -1 |
| GLY | 57.05 | 4.50 | 0 |
| HIS | 137.1 | 6.08 | 0.5 |
| ILE | 113.2 | 6.18 | 0 |
| LEU | 113.2 | 6.18 | 0 |
| LYS | 128.2 | 6.36 | 1 |
| MET | 131.2 | 6.18 | 0 |
| PHE | 147.2 | 6.36 | 0 |
| PRO | 97.12 | 5.56 | 0 |
| SER | 87.08 | 5.18 | 0 |
| THR | 101.1 | 5.62 | 0 |
| TRP | 186.2 | 6.78 | 0 |
| TYR | 163.2 | 6.46 | 0 |
| VAL | 99.07 | 5.86 | 0 |

The histone tails are modeled as fully flexible polymer chains (i.e., no energetic penalty for bending) with a bond energy modeled as in Eq. 2, with stiffness constant *k* = 10 kcal*/*mol*/*Å^2^ and equilibrium bond length *r*_0_ = 3.5 Å. Bonds are defined between adjacent amino acids along the protein backbone. The globular domains of the histones are regions which exhibit small structural fluctuations, and are largely *α*-helical. Using the 211-bp nucleosome reference structure, a Gaussian elastic Network Model (GNM) [14] is created by connecting all globular domain amino acid pairs that are closer than 7.5 Å, again using Eq. 2 where *k* = 10 kcal*/*mol*/*Å^2^ and *r*_0_ is the bond length in the reference structure.

We develop two versions of the model: (a) breathing (i.e., with nucleosomes that are allowed to breath spontaneously) and (b) non-breathing (i.e., with nucleosomes that are constrained to remain fully wrapped). In practice, these two versions differ in how the DNA beads are bound to the histone protein core. In the breathing case, DNA ellipsoids and amino acid beads interact exclusively via the electrostatic Eq. 4 and Lennard-Jones potentials (hydrophobicity potential with *λ*=1, Eq. 5) defined above—this leaves the DNA free to bind and unbind spontaneously due to thermal fluctuations (i.e., “breathe”), and slide around the nucleosome core. In contrast, non-breathing simulations further constrain nucleosomal DNA by permanently bonding it to the histone core using a GNM with the same 7.5 Å threshold and bond parameters provided above, preventing these nucleosomes from breathing and sliding, and hence, forcing them to remain fully wrapped.

To simulate the presence of an implicit solvent and allow sampling of the canonical ensemble, we use Langevin dynamics. We use the following LAMMPS fixes: `fix nve` and `fix langevin` (using the Gronbech-Jensen/Farago formulation [15]) for the amino acid particles, and `fix rigid/nve/small` and `fix langevin` with angular momentum for the DNA beads.

To create the initial 12-nucleosome chromatin structure, we replicate a single nucleosome and join these copies together following the DNA double-helix. Care must be taken to avoid steric clashes so we use short non-equilibrium MD runs to pull the two DNA end points on adjacent nucleosomes, in opposite directions before joining them; this creates the chromatin array in a linear conformation that resembles an extended ‘beads-on-a-string’ structure.

Equilibration of structures is achieved by slowing decreasing, in steps, the friction coefficient of the Langevin thermostat whilst increasing the timestep. Unless otherwise specified, all production runs are performed at 300 K with a Langevin damping period of 100000 fs and a timestep of 40 fs; these values are the largest that ensure simulation stability. It is important to note that the timescales in these coarse-grained simulations are not directly comparable to timescales in atomistic simulations. The units of femtoseconds here are implemented to achieve compatibility with the units of the potentials.

### D. Chemically-specific model parameters: Non-bonded potentials

#### 1. Hydrophobic interactions

The parameters for all amino acid pair interactions are taken from Ref. [9], which provide the Kim-Hummer model parameter set A for interactions involving globular domains, and the parameter set D for interactions involving only histone tails. The mass, *σ*, and charge are given in Table II. Cross terms for *σ*, for use in Eq. 5, are calculated using the combination rule *σ* = (*σ*_*i*_ + *σ*_*j*_)*/*2, where *σ*_*i*_ and *σ*_*j*_ are the individual *σ*s for the two amino acids involved in the interaction. The cutoff distance is set at *r*_*c*_ = 3*σ*. The values of *υ* and *λ* for each amino acid pair can be found in the Supporting Information of Ref. [9].

Due to the dominance of the electrostatic DNA self-repulsion at the salt conditions we explore in this work, we approximate the DNA–DNA pairwise hydrophobic interaction as zero: *σ*_DNA–DNA_ = 0, *λ*_DNA–DNA_ = 1, *υ*_DNA-DNA_ = 0, with mass and charge as given in Table III.

**TABLE III.** DNA parameters

| Particle | Mass (g/mol) | Charge ($e$) |
| --- | --- | --- |
| DNA ellipsoid | 650 | 0 |
| DNA phosphate | $10^{-6}$ | -1 |

To parameterize the hydrophobic interaction between DNA and the different amino acids—i.e., the parameters for the potential *E*_Hydrophobic_ between DNA and amino acids—we fit these parameters to optimize the DNA ellipsoid–amino-acid and the DNA phosphate–amino acid radial pair-wise distance distribution functions (RDF) of the coarse-grained simulations to match that computed from our 211-bp all-atom simulations of single nucleosomes. We compute the RDF as:

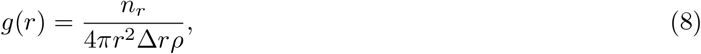

where *n*_*r*_ is the number of particles found at distance *r* in a spherical shell of thickness Δ*r*. To find *n*_*r*_ we construct a histogram of all pair distances, where *n*_*r*_ is the bin height and Δ*r* the bin width. *ρ* is the average density of the system. The fitted values are given in Table IV.

**TABLE IV.** DNA–amino acid interaction parameters.

| Interaction | $\sigma$ (Å) | $\epsilon$ (kcal/mol) | $\lambda$ |
| --- | --- | --- | --- |
| DNA ellipsoid–amino acid | 8 | 0.01 | 1 |
| DNA phosphate–amino acid | 4 | 0.1 | 1 |

#### 2. Electrostatic interactions

We set the relative permittivity as that for water *υ*_*r*_ = 80, to model the low concentration of monovalent ions within cells. The inverse Debye length *κ*, is varied with the monovalent salt concentration *c* (measured in units of mol/L) according to:

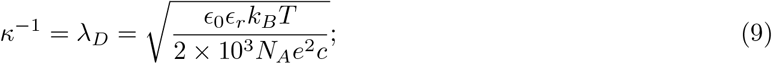

where *k*_*B*_ is the Boltzmann constant, *T* is the temperature, *N*_*A*_ is the Avogadro constant and *e* is the elementary charge. We set the cutoff distance, *r*_*c*_ = 3.5*λ*_*D*_.

## IV. MINIMAL CHROMATIN COARSE-GRAINED MODEL

### A. Energy function of minimal model

The total potential energy of the minimal chromatin model is

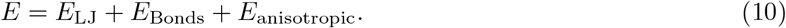

*E*_*LJ*_ is a Lennard-Jones interaction

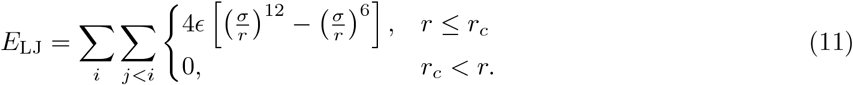

where *r* is the distance between the pair of interacting particles, *υ, σ* and *r*_*c*_ depend of the type of interacting particle. *E*_Bonds_ is similar to the RBP potential of the chemically-specific model, indeed it has the same functional form,

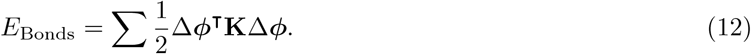

This RBP-like potential is instead parameterized to model DNA at a resolution of 1 bead per 5 base pairs. At this resolution the effects of DNA sequence become smoothed out, therefore all DNA bonds have the same values for **K** and ***φ***_0_.

The anisotropic term is a pairwise potential that depends on the relative orientations and shape of the interacting pair of ellipsoidal particles.

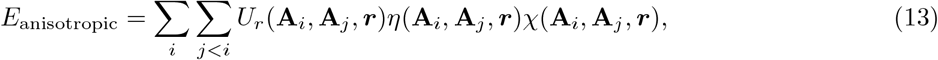

where **A**_*i*_ and **A**_*j*_ are the orientation matrices of particles *i* and *j* with center to center separation vector ***r***. This potential is a modified version of the well-know Gay–Berne potential [16] where we have replaced the Lennard-Jones like term with a cosine-squared term [17]. This allows for greater control over the depth and range of the potential. The *η* and *χ* terms are unchanged from the original version in LAMMPS [18].

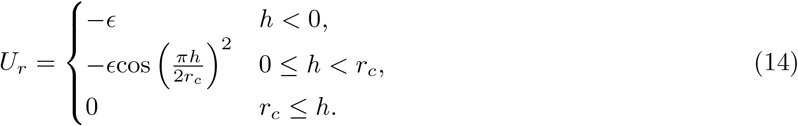

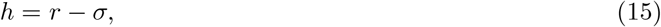

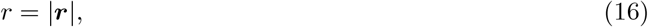

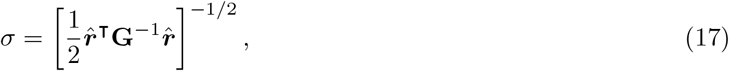

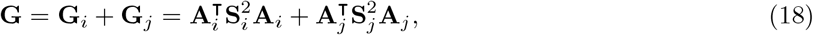

where **S**_*i*_ = diag(*a*_*i*_, *b*_*i*_, *c*_*i*_) is the shape matrix of particle *i* given by the ellipsoid radii and *r*_*c*_ is the cutoff distance (numerical value is potential specific). The *η* and *χ* terms contain the Gay-Berne parameters *µ,ν*, and the relative energy matrices 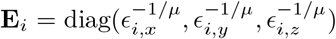 where *E*_*i,k*_ is the depth of the potential well in the direction of the *i*^th^ ellipsoids *k*^th^ axis. The anisotropic term only acts between the core particle and the DNA as is simulates DNA binding around the histone core.

The resulting potential of *E*_LJ_ and *E*_anisotropic_ acting on DNA beads due to the core bead is graphed in Figure S2.

**Figure S2.**
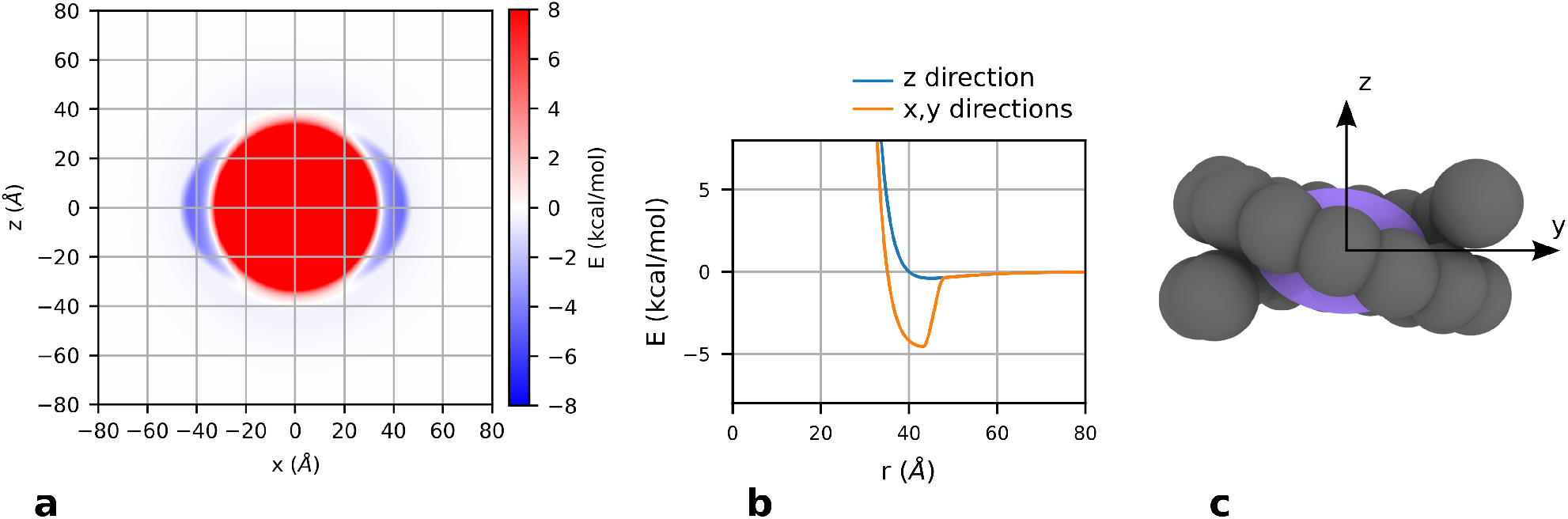
Description of minimal model anisotropic potential. **a**,**b** Graphs of the potential acting on the DNA beads due to the core bead; this includes both the LJ and anisotropic potential. The potential has a strong attractive region around the nucleosome *xy* plane and weak attraction at the *z* poles. It is symmetric around the *z* axis. **c** Illustration of DNA binding to the attractive region of the anisotropic potential.

### B. Definition and implementation of forces and torques

The forces and torques for the anisotropic potential are modified from the Gay–Berne implementation in LAMMPS [18, 19]. The force is

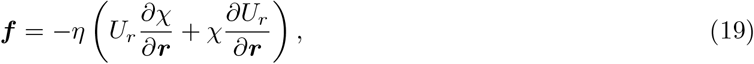

where

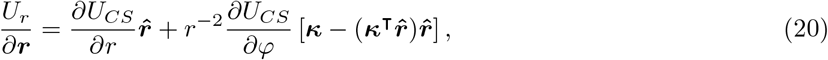

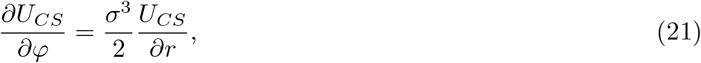

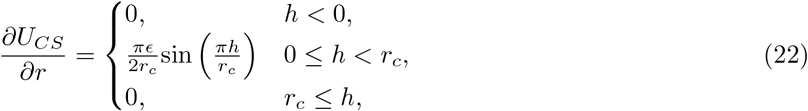

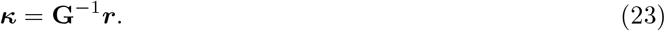

∂*χ/*∂***r*** is the same as in [18]. The torque on particle *i* is given by

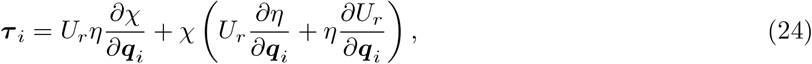

where

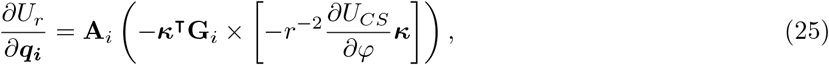

and ∂*χ/*∂***q***_*i*_ and ∂*η/*∂***q***_*i*_ are the same as in [18].

### C. Minimal model building & implementation

The minimal model uses two types of ellipsoidal particles. One represents the nucleosome core and is disk shaped (referred to as a core bead), the other type represents 5 DNA base pairs (referred to as a DNA bead) and is actually spherical but still uses the LAMMPS ellipsoidal type for computational compatibility reasons. The radii of the core beads is 28 *×* 28 *×* 20 Å and the radii of the DNA beads is 12 *×* 12 *×* 12 Å, these have been optimized as part of the fitting procedure. The beads are further categorized into linker and nucleosomal DNA, the nucleosomal DNA is fixed rigidly together with the core bead it is wrapped around to form a rigid nucleosome. The linker DNA beads are left free, bonded in sequence by the minimal RBP potential. The reference structures for creating the minimal model structures are the equilibrium structures from the chemically-specific model simulations (Figure S3). The minimal coarse-grained DNA beads are positioned by grouping every 5 lots of base-pairs and taking the average position and orientation. The minimal coarse-grained core beads are positioned by taking the center of mass of the nucleosome core globular domain. The orientation is set by using specific amino acid beads to consistently construct a nucleosome’s x,y,z orientation axis vectors.

The difference between the non-breathing and breathing in the minimal coarse-grained model is only the initial structures that are used, both have the DNA that is classified as nucleosomal rigidly fixed to the nucleosome core. For the breathing structures the effects of DNA unbinding/sliding is fixed into the structure from the equilibrated chemically-specific coarse-grained structure that was used as the reference. This means that although during a minimal coarse-grained simulation the DNA cannot further dynamically breathe, it is in configurations that represent the thermodynamic fluctuations that occur due to DNA breathing.

**Figure S3.**
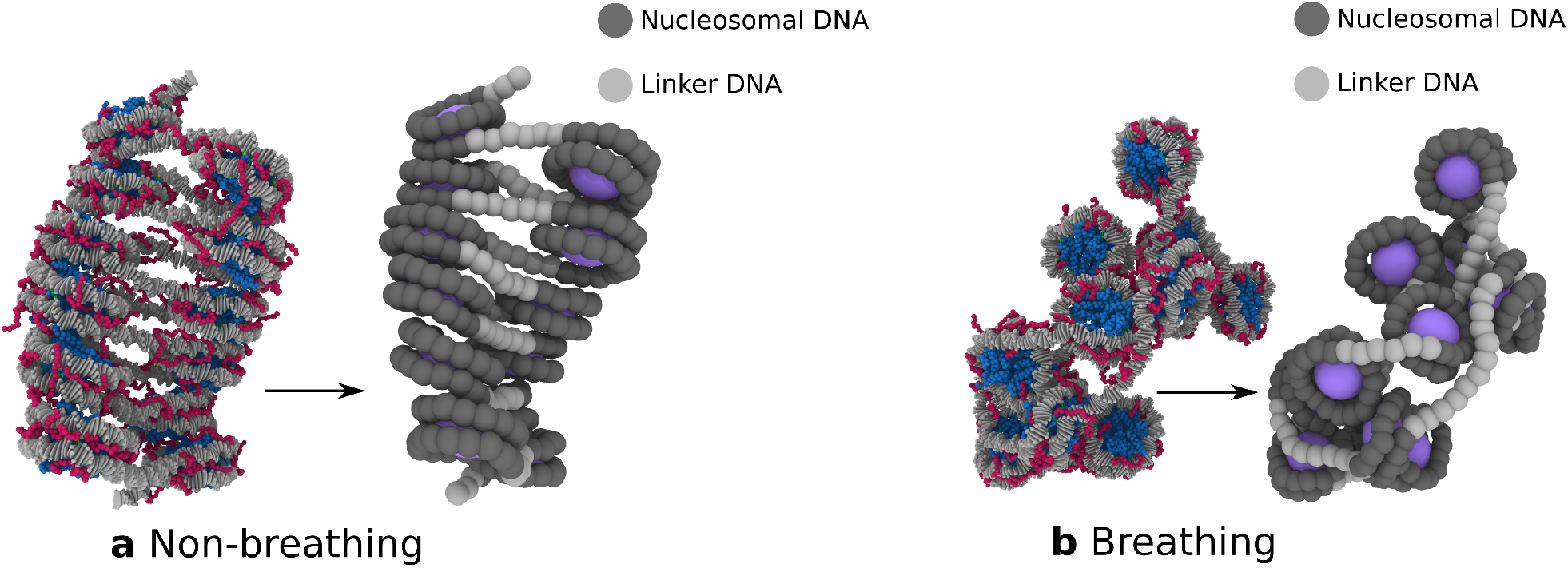
Mapping of minimal model initial structures from chemically-specific model equilibrium structures. The important difference between the non-breathing (**a**) and the breathing (**b**) chromatin is the amount of linker DNA, at higher salt the non-breathing chromatin has significantly more.

Once again Langevin dynamics is used, but for the minimal coarse-grained model the damping time constant is 5 ns and the timestep is 500 fs, the temperature is still 300 K. The particle masses are kept consistent with the chemically-specific model with the DNA beads having a mass of 3250 g/mol and the core beads having a mass of 100,000 g/mol.

### D. Fitting of the minimal chromatin model parameters from chemically-specific coarse-grained simulations

We fit the parameters of the minimal model such that is approximates the salt dependent behavior of the chemically-specific model.

To obtain initial values for Eq. 11 we first computed the inter-nucleosome PMFs using the chemically-specific model for high and low salt, these are shown in Figure S4 a1 and a2. The method used to calculate these is described in Section V D. We then performed a similar calculation using the minimal model. Due to the fact that the minimal nucleosomes are completely rigid, umbrella sampling is not needed, instead we simply compute the potential energy as a function of the inter-nucleosome distance. We optimize the values of *σ* and *υ* such that the minimal model best approximates the shapes of chemically-specific inter-nucleosome PMFs. Armed with these initial guesses of *σ* and *υ*, which are the end points of the salt range we wish to model, we proceed to find an adequate interpolation to model the salt dependent behavior. To do this we use the radius of gyration of non-breathing 12-nucleosome chromatin as the observable to compare between the minimal model and the chemically specific model. Using a combination of manual adjustment and grid search techniques we obtain the optimal parameters (in Table V) which give the *R*_*g*_ values in Figure S4 b2 which compare well with the chemically specific model.

**TABLE V.** Minimal model Lennard-Jones parameters. *c* is the salt concentration. *E*(*c*) and *S*(*C*) are linear interpolations of the data in table VI. Note that *σ*_*c*_ = 2^1*/*6^*σ* implies that the potentials are repulsive only.

| Interaction pair | $\epsilon$ (kcal/mol) | $\sigma$ (Å) | $\sigma_c$ (Å) |
| --- | --- | --- | --- |
| Core-core | 0.1 | 55 | $2^{1/6}\sigma$ |
| Core-DNA | $E(c)$ | 40 | $3\sigma$ |
| DNA-DNA | 0.1 | $S(c)$ | $2^{1/6}\sigma$ |

**TABLE VI.** *E*(*c*) and *S*(*c*) are found by linear interpolation of this data.

| $c$ (mol/L) | $E(c)$ (kcal/mol) | $S(c)$ (Å) |
| --- | --- | --- |
| 0.15 | 0.4 | 24 |
| 0.115 | 0.375 | 24.5 |
| 0.082 | 0.35 | 25 |
| 0.07 | 0.3 | 26 |
| 0.06 | 0.25 | 27 |
| 0.052 | 0.2 | 28 |
| 0.05 | 0.1 | 30 |
| 0.042 | 0.01 | 34 |

**Figure S4.**
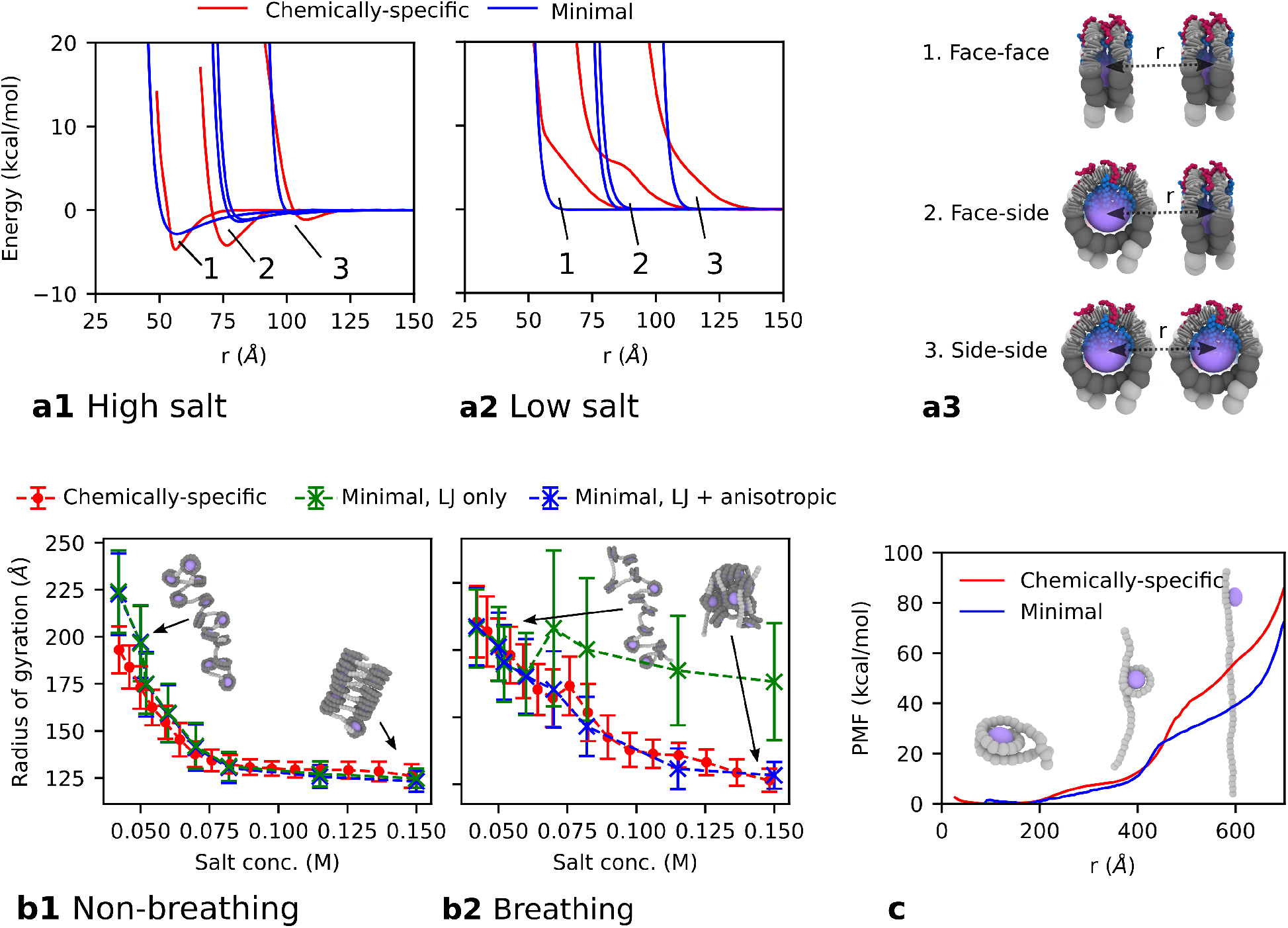
Fitting of the minimal model potentials and parameters to the behavior of chromatin within the chemically-specific model. **a** Orientational-dependent interactions between two nucleosomes at high (0.15 M) and low (0.05 M) concentration of NaCl, **a1** and **a2** respectively, for the chemically specific (red) and minimal (blue) models. The chemically-specific model curves are PMFs computed using the center-to-center nucleosome distance among nucleosome pairs in the three different relative orientations described in **a3**. The minimal model curves represent the total pairwise energy as a function of center-to-center nucleosome distance in the three same orientations. **b** Salt-dependent radius of gyration of 12-nucleosome 165-bp chromatin for the chemically–specific model and the minimal model, the error bars are the standard deviation. The figure compares the values computed with and without the anisotropic potential for chromatin with non-breathing **b1** and breathing **b2** nucleosomes. This illustrates the need of the anisotropic potential to recover the behavior of chromatin with breathing nucleosomes at higher salt. **c** Nucleosome unwrapping PMFs for both models showing good agreement in the low extension regime.

Moving on to the breathing model and comparing the radii of gyration between the minimal model and the chemically-specific model (red and green lines in Figure S4 b2), we find a significant difference in the behavior at higher salt values. This is due to the unwrapped DNA not having strong enough interactions to the exposed nucleosome cores. To account for this we developed the anisotropic potential which provides a strong short range attractive binding potential around the nucleosome core where the DNA is located in an archetypal nucleosome.

To fit the parameters of the aniostropic potential we first use our knowledge of the shape of the nucleosome core and the region where the DNA binds. This allows us to set the values of the ellipsoid shape matrices **S**_core_ and **S**_DNA_, and the anisotropic energy matrices **E**_core_ and **E**_DNA_. Specifically **E**_core_ is constructed such that there is no attraction at the z-poles of the ellipsoid. We then ensure that it is short ranged enough to have no effect on the non-breathing minimal model. That is, the potential well is completely covered by the bound nucleosomal DNA. This gives us the value of cutoff *r*_*c*_, and the correspondence of the green and blue points in Figure S4b1 demonstrates the value ensures the anisotropic potential has no effect on the non-breathing minimal model. To fit the remaining parameter *E*, the depth of the potential, we compute the PMF of nucleosome unwrapping for a minimal model nucleosome where the DNA is completely free from the core and only the pairwise interactions keep it bound. We perform a grid search parameter sweep and chose the value of 6 kcal/mol which gives the PMF that best approximates the chemically-specific PMF in the low extension regime. The resulting PMF is shown in Figure S4c.

Finally, the minimal RBP-like helical parameters are optimized by directly fitting to chemically-specific coarse-grained simulations of 200bp DNA strands. 10 simulations of random DNA sequences are performed for 10 million timesteps. The chemically-specific coarse-grained trajectories are then mapped into the minimal representation. For each timestep and each bond, the helical parameters between the minimal ellipsoids are computed. We now have histograms of the equilibrium distributions for each of the six helical parameters. We observe that shift, slide, tilt, and roll are centered about roughly zero, so we set these equilibrium values to zero. Rise and twist are set to the calculated mean values. The stiffness matrix is constructed by computing the variance of each helical parameter.

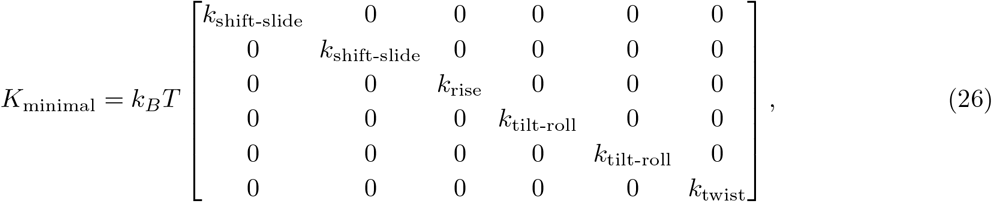

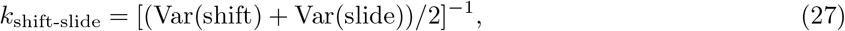

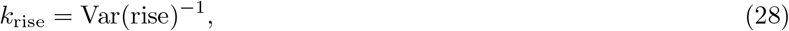

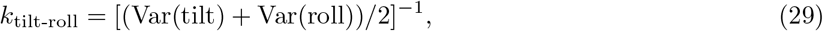

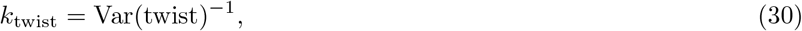

where Var represents taking the variance. Note that here, unlike the original base-pair resolution RBP model, we neglect the off diagonal covariance terms. Furthermore, shift and slide are set equal, and tilt and roll are set equal. At this coarse-grained level we are unable to resolve the DNA major/minor grooves so the approximation of symmetry around the z-axis is acceptable. These approximations are an important step as it allows us to redefine the equilibrium value of twist to be zero and remap the minimal DNA in the structures by rotating each DNA bead by:

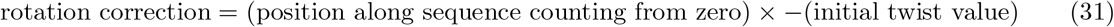

This is needed because the initial twist value of 167^°^ causes numerical instabilities in the current computational implementation of the RBP model due to the angles wrapping around from 180^°^ to -180^°^. We confirm that the DNA persistence length of the minimal model is the similar to that of the chemically-specific model.

### E. Parameters of the minimal model potentials

The Lennard-Jones interaction parameters are in table V, some are dependent on the salt concentration *c*.

The RBP-like bonds are have the following parameters:

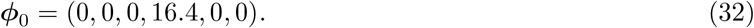

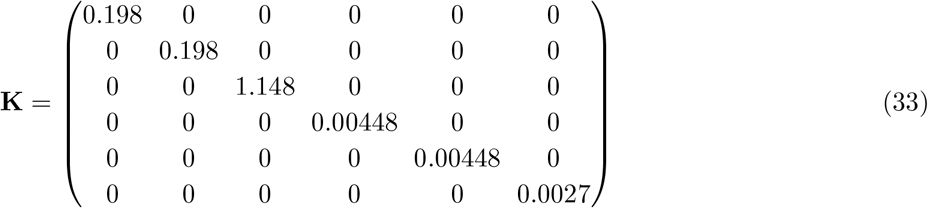

where the units are Å for distances, degrees for angles, kcal/mol/Å^2^ for spatial stiffness matrix elements, and kcal/mol/degrees^2^ for angle stiffness matrix elements.

The parameters for the anisotropic potential are in table VII.

**TABLE VII.** Anisotropic potential parameters. diag(a,b,c) means diagonal 3×3 matrix with the elements a,b,c on the diagonal.

| Parameter | Value |
| --- | --- |
| $\mathbf{S}_{\text{Core}}$ | diag(28, 28, 20) |
| $\mathbf{S}_{\text{DNA}}$ | diag(12, 12, 12) |
| $\epsilon$ | 6 kcal/mol |
| $r_c$ | 5 Å |
| $\mathbf{E}_{\text{core}}$ | diag($1^{-1/\mu}$, $1^{-1/\mu}$, $0.0001^{-1/\mu}$) |
| $\mathbf{E}_{\text{DNA}}$ | diag($1^{-1/\mu}$, $1^{-1/\mu}$, $1^{-1/\mu}$) |
| $\mu$ | 1 |
| $\nu$ | 1 |

## V. SIMULATION PROTOCOLS AND ALGORITHMS

### A. Debye-length replica exchange simulation method

We implement our Debye length Hamiltonian Replica Exchange Molecular Dynamics (HREMD) method via a modified version of the existing LAMMPS parallel tempering command from the `REPLICA` package.

Simultaneous simulations are performed on different replicas of the system, with the implicit salt concentration varying (by changing the Debye length) between replicas. Each time a Hamiltonian exchange between two replicas is attempted, the potential energies of the replicas are calculated as if they have swapped Debye lengths. The exchange probability is then determined using the Metropolis criteria,

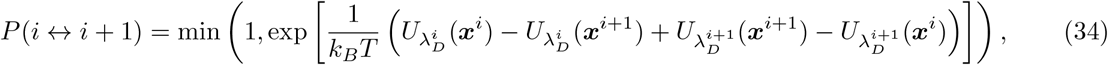

where ***x***^*I*^ is the chromatin coordinates of the *i*^th^ replica and 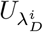 the potential energy function at Debye length 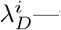 —the original Debye length of the *i*^th^ replica. Unlike temperature replica exchange, HREMD requires the recalculation of the potential energy. However, here only the re-computation of *E*_Electrostatic_ is necessary, and as a replica exchange attempt is only made infrequently this does not significantly degrade performance.

### B. Details of chemically-specific model 12-nucleosome chromatin simulations

HREMD simulations with 16 replicas with Debye lengths ranging from 8–15 Å (see Table VIII) are performed for 12-nucleosome chromatin arrays with NRLs of 165 bp and 195 bp in both the breathing and non-breathing cases. The simulations were run for ∼100 million timesteps and a set of exchanges were attempted every 10,000 timesteps. Each set of exchanges either attempts to exchange replicas {1-2,3-4…15-16} or {2-3,4-5…14-15}, with each set picked with a 50% probability.

**TABLE VIII.** HREMD Debye Length (*λ*_*D*_) values.

| Replica | 1 | 2 | 3 | 4 | 5 | 6 | 7 | 8 | 9 | 10 | 11 | 12 | 13 | 14 | 15 | 16 |
| --- | --- | --- | --- | --- | --- | --- | --- | --- | --- | --- | --- | --- | --- | --- | --- | --- |
| $\lambda_D$ (Å) | 8.00 | 8.34 | 8.70 | 9.07 | 9.46 | 9.86 | 10.29 | 10.73 | 11.19 | 11.67 | 12.16 | 12.69 | 13.23 | 13.79 | 14.38 | 15.00 |

For qualitative assessment of chromatin dynamics an equilibrium structure was taken from the 0.1 mol/L simulations for both the breathing and non-breathing simulations, and run using standard MD (non replica exchange) for ∼40 million timesteps. They were used for the generation of the *k* = 2 interaction time series plotted in main text Figure 5.

### C. Details of the chemically-specific model nucleosome unwrapping potential of mean force simulations

The force–extension simulations of mononucleosomes were performed following the umbrella sampling procedure proposed by Lequieu et al. [20] but with a tension of zero. The collective variable is the DNA extension which is the distance between the first and last base-pair. We implement the umbrealla sampling simulations using the COLVARS library in LAMMPS [21] (version 2019-08-05). Starting from an equilibrium structure with an extension of 25 Å, initial configurations for the windows were prepared via constant velocity steered MD (SMD) until the extension was at 750 Å. A spring constant of 0.01 kcal/mol/ Å^2^ was used with a pulling velocity of 9.0 *×* 10^−6^ Å/fs, giving a total pulling time of 100 ns. The extension range was split into 50 equally spaced windows. Each window was run with a fixed harmonic biasing potential at the corresponding extension with a spring constant of 0.025 kcal/mol/ Å^2^ for 100 ns. These values were chosen by assessing histogram overlap and checking that the calculated potential of mean forces (PMF) were the same on longer timescales. The entire procedure was repeated 5 times and the aggregate data was used for computing the PMF via the Weighted Histogram Analysis Method (WHAM) [22]. The same procedure was done for all nucleosome configurations and environments listed in Figure S8.

### D. Details of the chemically-specific model nucleosome-nucleosome potential of mean force simulations

Due to the disk like shape of nucleosomes the interaction is divided into three orientation dependent configurations: face-face, face-side, and side-side as illustrated in Figure S9. A PMF is constructed for each configuration using umbrella sampling as follows. An initial single nucleosome structure is equilibrated, it is then replicated and each copy is positioned such that the center to center distance is 200 Å. The nucleosomes are rotated into the correct orientations and held fixed using COLVARS’ [21] `angleOrient` collective variable with a harmonic restraint of strength 1 kcal/mol/degrees^2^ with center 0^°^. A further restrained collective variable `distanceDir` is used to keep the nucleosomes aligned along the z-axis. The harmonic restraint center is the (0,0,1) unit vector and the force constant is 10 kcal/mol. The final collective variable is distance between the nucleosome centers, *R*.

To prepare initial configurations for the windows, *R* is varied using a SMD protocol from its initial value of 200 Å to its final value of 10 Å over 1 million timesteps with a force constant of 0.1 kcal/mol. The range of *R* was split into 39 equally spaced windows, from 10 to 200, and each window was run for 10 million timesteps at the corresponding value of *R* with a force constant of 0.05 kcal/mol/ Å^2^. The orientational collective variables are kept with the same restraints as in the setup stage. Finally the PMFs were computed from the window trajectories using WHAM [22].

### E. Estimation of radius of gyration from simulations with our minimal model

For the radius of gyration simulations plotted in Figure S4 we first obtain 50 random initial 12-nucleosome structures by generating them as described in Section IV C from randomly selected, equilibriated, structures from the chemically-specific model HREMD simulations at the corresponding salt. We then run the 50 minimal model 12-nucleosome structures for 500,000 timesteps. These 50 repeats are done for each plotted salt value.

### F. Nucleosome unwrapping from potential of mean force simulations with our minimal model

To calculate the PMF plotted in Figure S4 we once again use the COLVARS library and follow a very similar procedure to Section V C. The distance between the first and last DNA bead *R*, is varied from 100 Å to 700 Å in 60 steps, with each step lasting 10,000 timesteps. The time series of *R* is split into the corresponding 60 windows and WHAM [22] is used to calculate the PMF.

### G. Direct coexistence simulations

In order to compute the phase diagram of 12-nucleosome chromatin we employ the direct coexistence method [23–25] using 125 independent 12-nucleosome chromatin arrays with 165-bp NRL at different conditions. The chromatin array initial structures are obtained by generating them from randomly selected, equilibrated, structures from the chemically-specific model HREMD simulations at the corresponding salt.

In a direct coexistence simulation one places both phases; i.e., the dilute liquid and condensed liquid phase in the same box. The simulation is performed until they reach equilibrium at their coexistence densities. Once equilibrium is reached to measure the density of coexistence we compute an average density profile along the long side of the simulation box with the center of mass fixed. The density profiles for our simulations are shown in Figure S5.

We estimate the critical salt concentration *c*_*c*_ by fitting the density difference between the coexisting low-density *ρ*_*l*_(*c*) and high-density *ρ*_*h*_(*c*) phases to the expression [26],

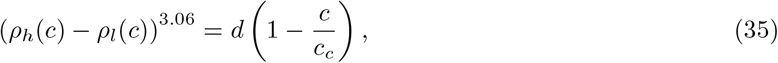

where *d* is a fitting parameter. The critical density *ρ*_*c*_ is estimated using the law of rectilinear diameter,

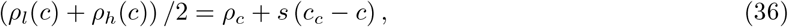

**Figure S5.**
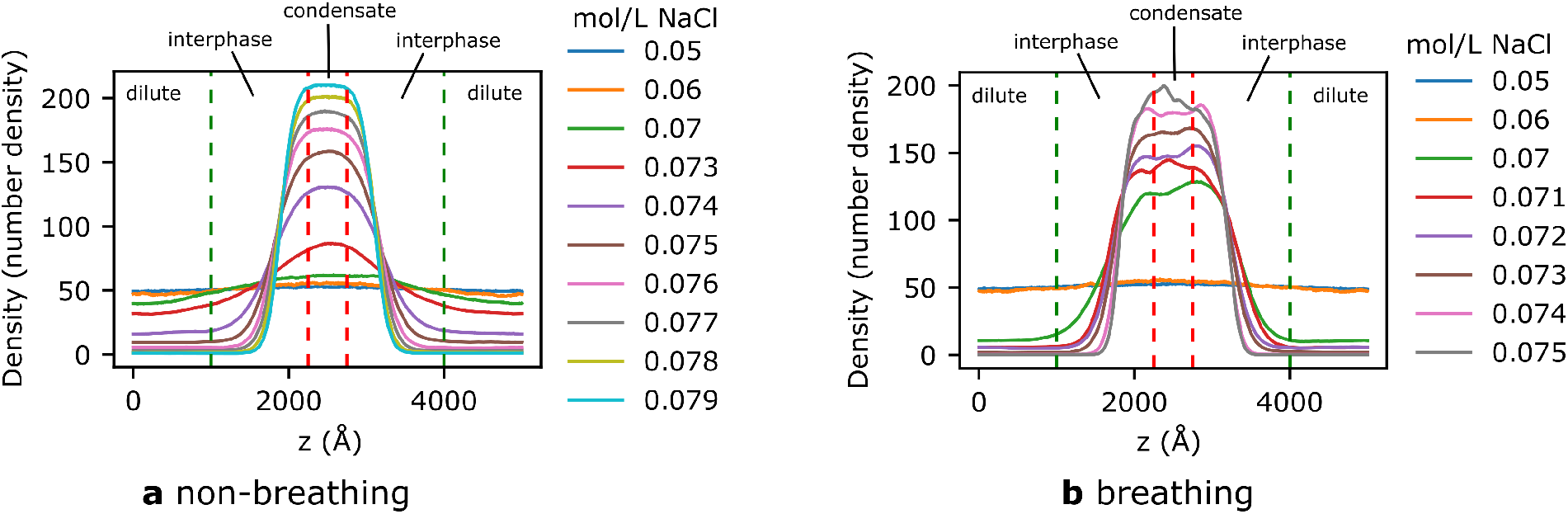
Equilibration of simulations and determination of chromatin density. Density profiles along the box long axis z, of a phase-separated system obtained via direct coexistence simulations at varying salt concentrations for chromatin with (**a**) non-breathing and (**b**) breathing nucleosomes. The presence of sharp, non-fluctuating interfaces (the two regions between the green dashed lines and the red dashed lines) is indicative of phase separation. The density of the condensate (middle region enclosed by the two red dashed lines) and dilute phase (outer regions from the extremes of the box to the green dashed lines) are averaged to determine the respective chromatin densities. The interface regions are excluded from these calculations.

where s is a fitting parameter.

## VI. FURTHER DETAILS ON CHEMICALLY-SPECIFIC MODEL VALIDATION

### A. Nucleosome formation

We investigate the ability of the chemically specific model to spontaneously form nucleosomes starting from configurations where the DNA is completely unwrapped and unbound from the histone core. These simulations are performed by setting up the structures, as pictured in the left panel of Figure S6a, in a periodic simulation box. The box dimensions are approximately double the length of the fully extended DNA, the periodicity ensures the DNA and nucleosome will eventually come in to close enough contact so they can interact. The simulations are run until binding occurs and then run for a further 10 million timesteps to ensure the resulting structures are well equilibrated. 64 trials were run and categorized by visual inspection into the pie chart categories in Figure S6a. We observe that a nucleosome is only correctly formed approximately one third of the time, the other resulting configurations include: a reversome, which is when the DNA is wrapped in a right-handed helix; “knotted” configurations where the DNA coiling is nucleosome-like but has DNA overlap; finally “wrong” where the DNA coiling is not nucleosome-like. This suggests additional processes are required to ensure the model always forms nucleosomes. Experimental literature shows that with completely relaxed DNA the formation of nucleosomes is significantly slowed, with positive supercoiled DNA nucleosomes do not form, and with negative supercoiled DNA the nucleosomes form instantaneously [27]. To investigate the response of the model to DNA supercoiling we repeat the simulations with the addition of torsional restraints to the DNA strand which is setup with either negative or positive supercoiling. The supercoiling is achieved by either increasing (for positive) or decreasing (for negative) the twist angle between base-pairs by 10%. The torsional restraints fix the rotation of the end-two base-pair by the addition of strong additional forces such that they cannot rotate about their z-axis. Additionally we start the simulation with the DNA in contact with the histone core. Performing 64 repeats we observe that negative super-coiling always forms a nucleosome, consistent with the experimental literature, and positive supercoiling always forms the chirally inverted reversome (Figure S6b). The reversome is a metastable state of a nucleosome that is observed in experiments [28–30].

**Figure S6.**
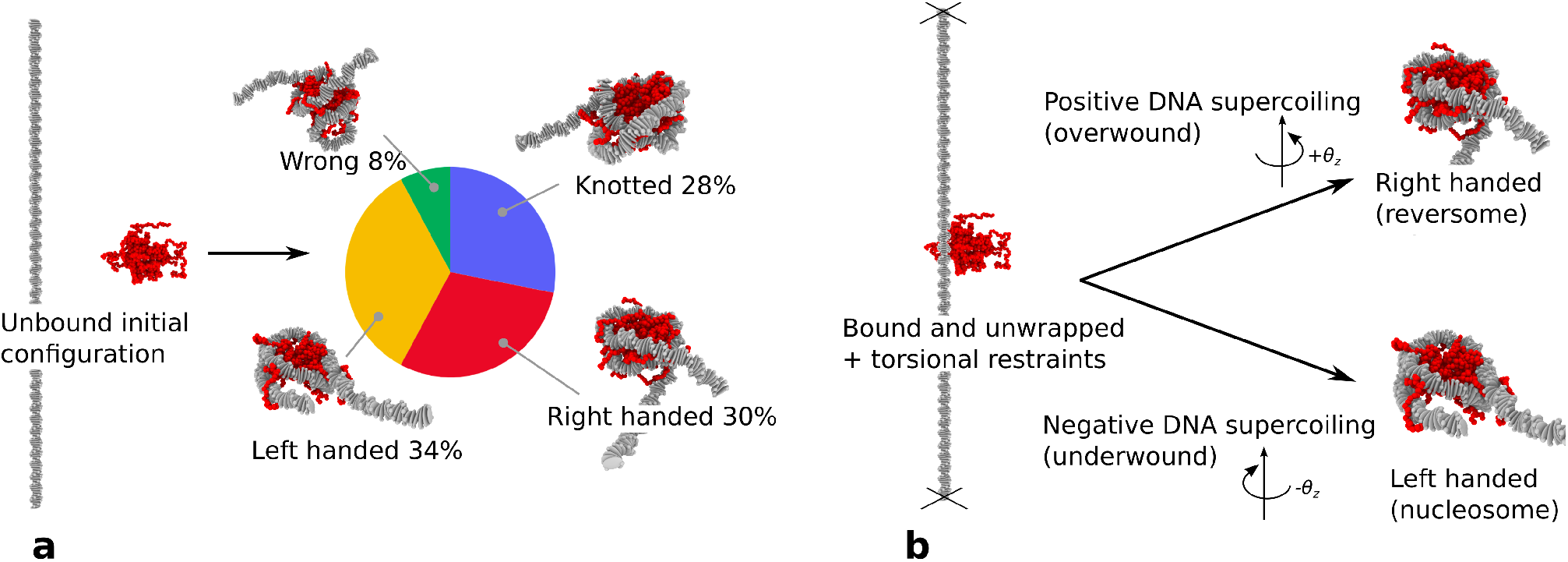
Spontaneous formation of nucleosomes predicted by our chemically-specific coarse-grained model at 0.15 mol/L NaCl and 300 K. **a** Starting from an unbound DNA to histone core configuration, i.e., where the DNA and histone octamer are completely separated, as time progresses, the DNA wraps around the histone core in one of four possible ways: left-handed supercoiling (a canonical nucleosome) with a ∼34% probability, right-handed supercoiling (a reversesome) with a ∼30% probability, a knotted configuration which is neither a left-handed or right-handed supercoil but still resembles a nucleosome-like shape with a ∼28% probability, and other configurations which differ significantly from a nucleosome topology (“wrong”) with a 8% probability. **b** Starting with a fully unwrapped nucleosome, but with the histone core bound to the DNA at a single point, we apply a weak super-coiling. In this case, as time progresses, the model always form nucleosomes when DNA is underwound, and reversomes when DNA is overwound.

### B. Persistence length of DNA

To test our chemically-specific coarse-grained DNA model we compute the persistence length of DNA as a function of monovalent salt concentration (Figure S7a) and DNA sequence (Figure S7b). The simulations are performed using 300 bp long strands of isolated DNA. 10 repeats are done for each data point for a total simulation time of ∼100 million time steps. The persistence length *P* of polymer is the length at which correlations on the direction of the polymer tangent are lost. We use the following definition:

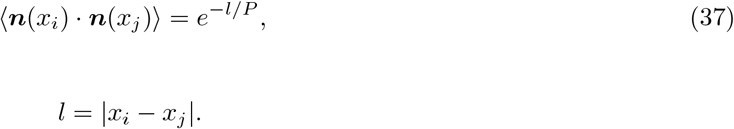

Where ***n***(*x*_*i*_) is the tangent vector of the polymer at location *x*_*i*_—note that location here is in terms of contour distance with units of base-pairs (symbol bp). E.g., *x*_1_ = 1 is the 1st base-pair and *x*_10_ = 10 is the 10th base-pair. *l* is the contour distance. To compute the value of *P* from our simulations we use the DNA ellipsoids’ quaternion orientation to directly give the tangent vectors. The end 20 base-pairs are excluded from analysis. For each time step all pairs of ***n***(*x*_*i*_) ·***n***(*x*_*j*_) are computed and the average is taken over all timesteps for each *l* value. We then plot the scatter graph of *l* verses ⟨***n***(*x*_*i*_) ·***n***(*x*_*j*_) ⟩ and fit an exponential curve to obtain *P* .

We see reasonable agreement with the experimental data for the salt dependence, the differences between our values and the experimental are of the same order as the differences between the the two experimental plots. This highlights the difficulty in accurate measurement of the persistence length of semi-flexible polymers with the existence of multiple techniques and alternative definitions to our Eq. 37 [35–37]. For the sequence dependent behavior our results follow the general trend where stiffness decreases going from left to right across the sequences. All but one data point is either in agreement with the experimental or the MD results. There are significant differences between all the different plots which can be explained by the aforementioned difficulty in accurate persistence length calculations.

**Figure S7.**
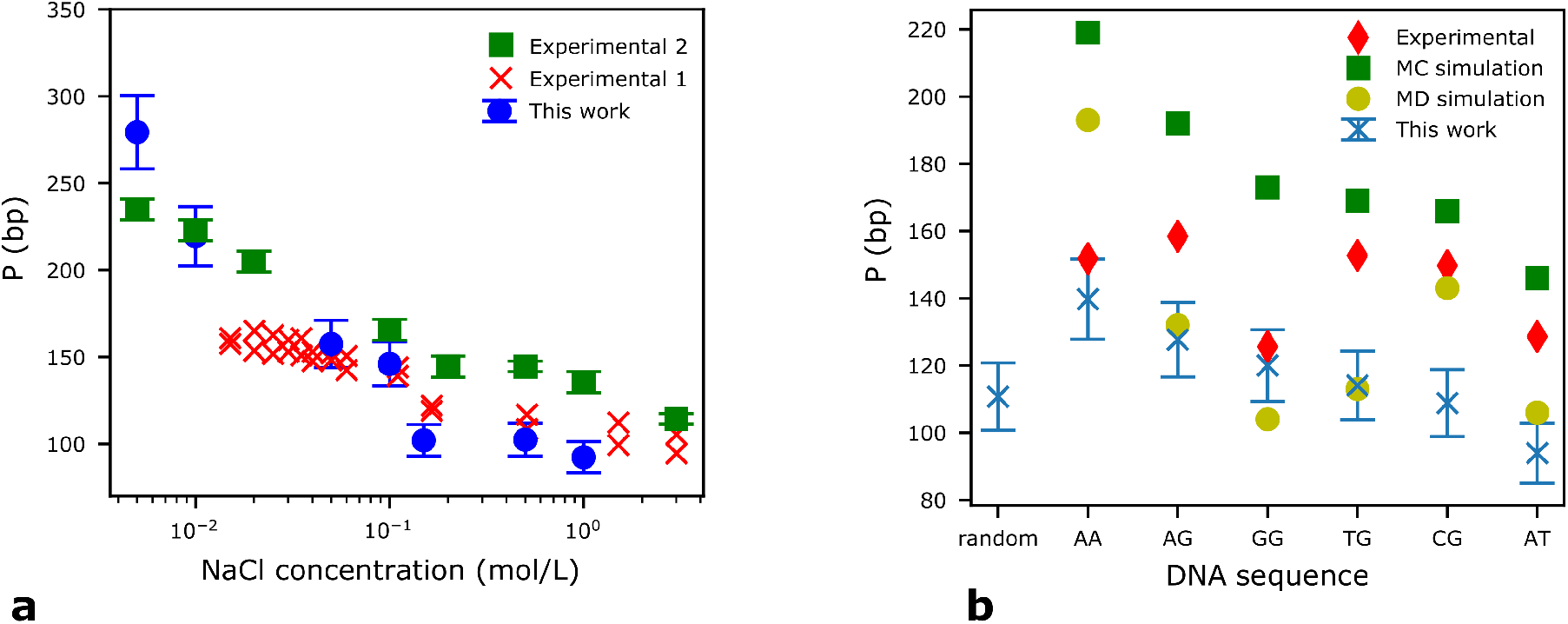
Experimental validation of the modified DNA rigid base pair model with added phosphate charges. **a** Persistence length of DNA in units of base-pairs (bp) as a function of NaCl concentration in mol/L at 300K. Blue circles are the values obtained with our simulations on a set of 300 bp DNA strands with varying sequences. Red crosses (experimental 1) are values from single-molecule high-throughput tethered particle motion experiments on DNAs of 1201 and 2060 bp at room temperature [31]. Green squares (experimental 2) are values from Rayleigh light scattering experiments for a T7 bacteriophage DNA from [32]. **b** Persistence length of DNA as function of DNA sequence from our simulations (blue crosses) using a random sequence and six poly(XY) sequences for DNA of 300 bp in length. We compare to values from experimental cyclization assays [33], coarse-grain Monte Carlo simulations [34], and all-atom MD simulations [34].

### C. Comparison with force-spectroscopy experiments of mononucleosomes

The force-extension curves in Figure S8a are computed as the numerical derivative of the PMF curves in figure 2a of the main text. The rupture forces listed in Figure 2c are the values at the peaks, labeled as *F*_1_ and *F*_2_ in Figure S8a, that occur at the transition states.

During the first regime (state 1), the force increases moderately as the DNA extends until it reaches a local maximum. The force then drops signaling the DNA-protein interactions being broken as the outer DNA turn unwraps. In the second regime (state 2), a similar behavior occurs with the force increasing significantly as the DNA continues to extend, and then reaching a second maximum and dropping again. This second drop in the force corresponds to the rupture of the DNA-protein interactions in the inner DNA turn. The third and final regime (stage 3) reflects the stretching of the fully unwrapped, but still histone-bound, nucleosomal DNA.

The forces required to unwrap the outer and inner DNA turns according to our model—e.g., ∼5 pN and ∼30 pN at 0.15 mol/L for the outer and inner DNA turn, respectively—are in accordance with the values from force spectroscopy experiments [38–40].

### D. Force-extension simulations of oligonucleosomes

As further validation, we conducted force-extension simulations of 4-nucleosome chromatin arrays by performing constant velocity steered MD. The spring constant was 0.001 kcal/mol/Å^2^ and the pulling velocity was 3.0 *×* 10^−6^ Å/fs The initial structures were equilibrated for 100 ns and then the SMD procedure was run for 1000 ns. Values of the applied SMD force and current extension are stored at each timestep (Figure S8b).

In Figure S8b, we compare the force-extension response of a 4-nucleosome system using our breathing model and the non-breathing model. Our force-extension curves exhibit the typical saw-toothed pattern observed in optical tweezer experiments of chromatin [41]; that is, the force exhibits an abrupt drop accompanied by a certain increase in the extension due to the partial unwrapping of individual nucleosomes. When we use the non-breathing model where the nucleosomal DNA remain permanently bound to the histone core the pattern disappears, and a much stiffer chromatin emerges.

**Figure S8.**
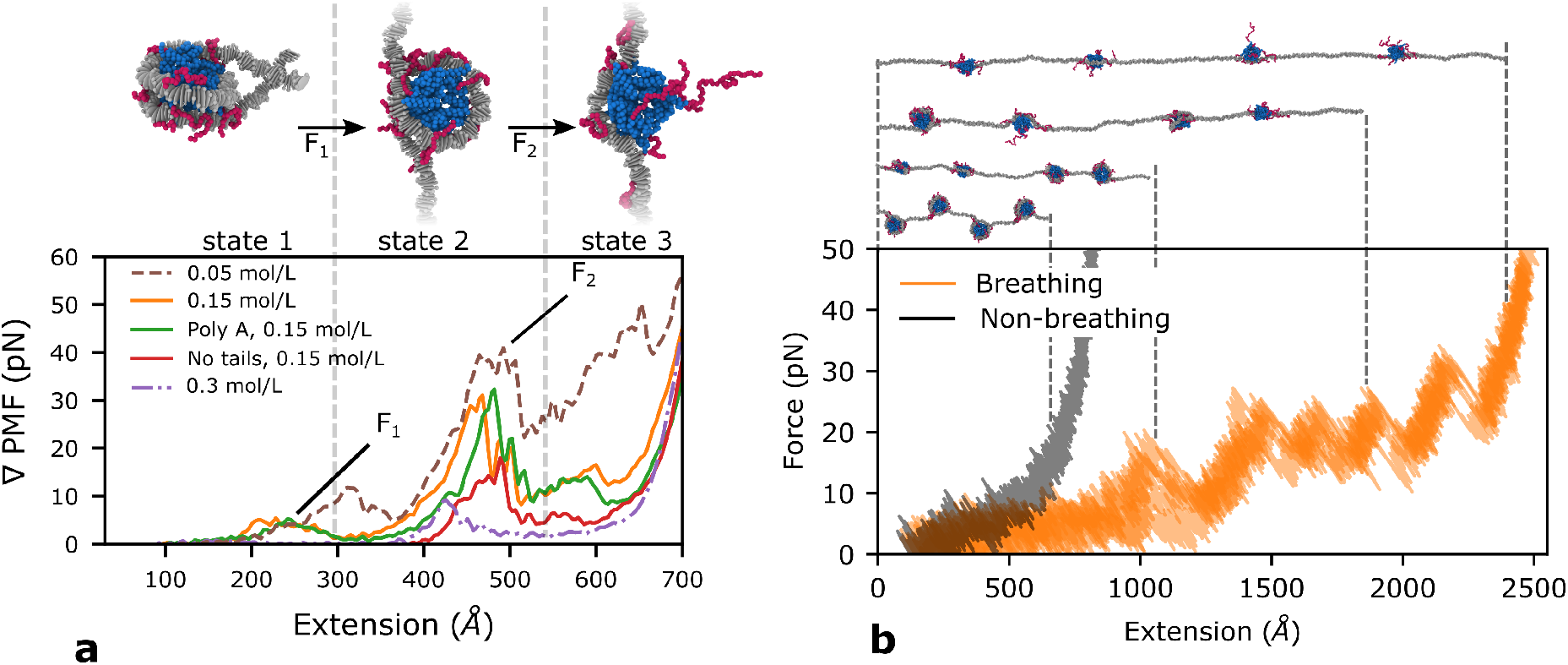
Force-induced unwrapping behavior of mononucleosomes and chromatin under varying conditions with our chemically-specific coarse-grained model. **a** Model predictions for the force-induced unwrapping behavior of mononucleosomes under varying conditions. Top: Representative simulation snapshots of nucleosome configurations (color coded as in Figure 1 of the main–Level 2) at three different stages of the unwrapping process, showing a fully wrapped nucleosome (state 1) at low pulling forces (≤*F*_1_), a nucleosome with the first turn unwrapped (state 2) at intermediate forces (*F*_1_–*F*_2_), and a fully unwrapped nucleosome (state 3) at higher forces (≥ *F*_2_). Bottom: Force (computed from the gradient of the PMF) in pN for nucleosome unwrapping as a function of the end-to-end DNA distance (or extension). The dashed brown, solid orange, and dashed purple curves correspond to 601-nucleosome simulations at 0.05 M, 0.15 M, and 0.3 M of NaCl, respectively. The green curve corresponds to simulations of a poly-A nucleosome at 0.15 M NaCl. The red curve was calculated for a nucleosome with all histone tails clipped at 0.15 M NaCl. **b** Force-extension for 4 nucleosome chromatin. There are 5 curves overlaid for chromatin with breathing nucleosomes (orange) and one curve using the fixed non-breathing version of the model for reference (black). Illustrated conformations are shown at the top with the extension indicated by the dashed lines.

## VII. ADDITIONAL ALGORITHMS

### A. Quaternion to rotation matrix

The unit quaternion

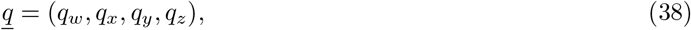

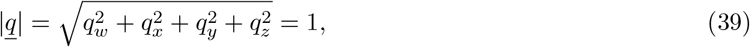

can be converted into an orthogonal rotation matrix:

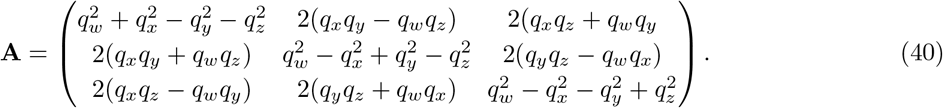

### B. Computation of helical parameters

We compute the helical parameters of the RBP model using the SCHNAaP procedure [8]. The two base-pairs (DNA ellipsoids) have positions and orientation quaternions ***r***_1_, *<u>q</u>*_1_ and ***r***_2_, *<u>q</u>*_2_ respectively. The following method computes the helical parameters ***φ*** = (*D*_*x*_, *D*_*y*_, *D*_*z*_, *τ, ρ*, Ω) that describe their relative orientation:

1. Convert quaternions *q*_1_ and *q*_2_ to matrices **T**_1_ and **T**_2_ whose columns are the 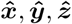, axis direction vectors.
2. Calculate the roll-tilt angle Γ:

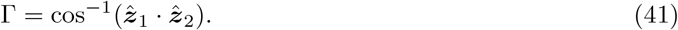
3. Calculate the roll-tilt axis ***rt***:

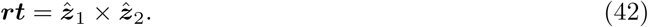
4. Rotate base-pair 1 and 2 about ***rt*** by +Γ*/*2 and −Γ*/*2 respectively:

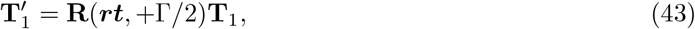

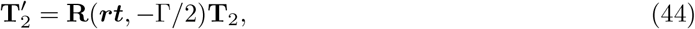

where **R**(***a***, *θ*) is an orthogonal matrix that describes a rotation of *θ* about axis ***a***
5. The mid-step matrix **T** is the mean of the rotated matrices:

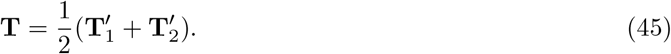
6. Twist Ω is the angle between the transformed y axis (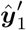 is the second column of 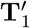):

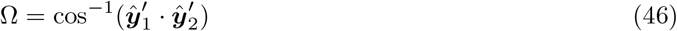
7. Calculate *φ* the angle between the roll-tilt axis and the mid-step y-axis (***ŷ***_*ms*_ is the second column of **T**):

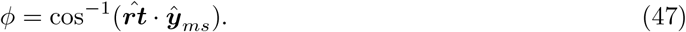
8. Roll *ρ* and tilt *τ* are given by:

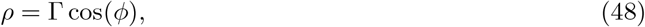

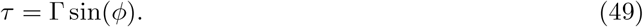
9. Shift *Dx*, slide *Dy*, and rise *Dz* are calculated as:

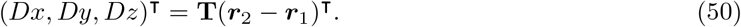

## VIII. ANALYSIS OF SIMULATION DATA

### A. Analysis of chemically-accurate coarse-grained simulations

#### 1. Orientation dependent nucleosome-nucleosome interactions

The relative orientation of two nucleosomes can be categorized into three states: face-face (ff), face-side (fs), and side-side (ss) as illustrated in Figure S9. To characterize these, we compute the nucleosome orientation matrices—the columns of which are the orthogonal unit axis vectors of the nucleosome. The center of a nucleosome is defined as the center of mass of the globular domain beads. The x axis passes through the nucleosome dyad and the z axis points perpendicularly out of the nucleosome “face” as shown in Figure S9.

**Figure S9.**
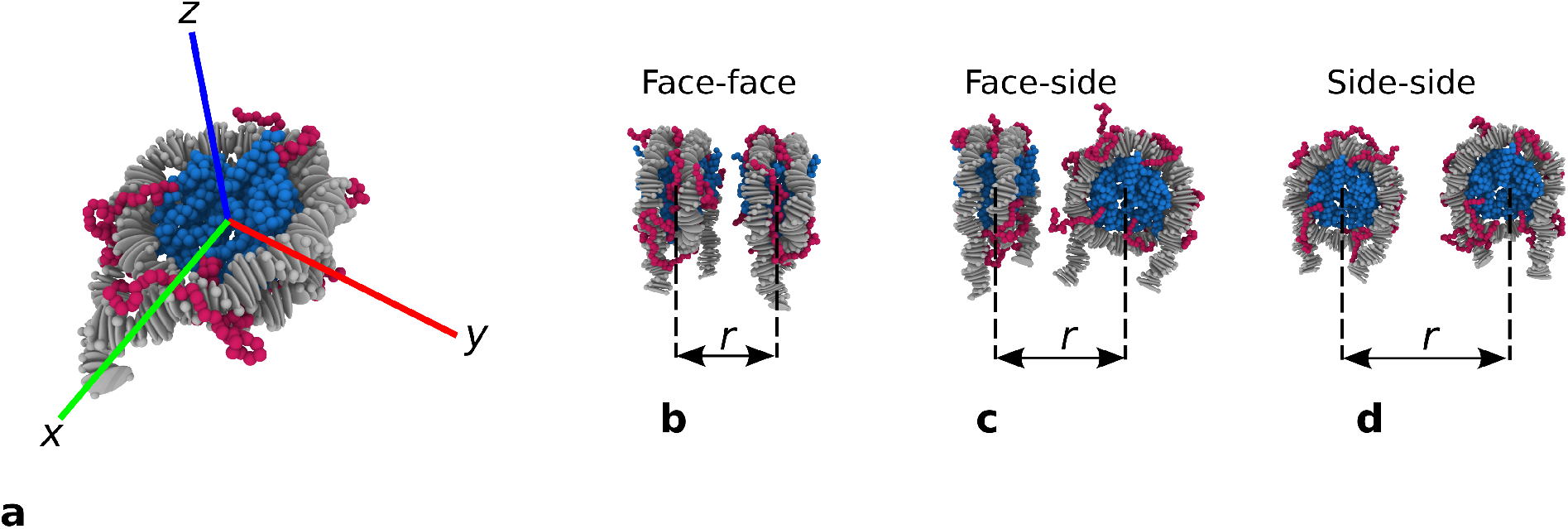
Definition of nucleosome pair orientations. **a** Nucleosome orientation axis: *x* points from the center of the nucleosome to the dyad position, *z* points out of the top face, and *y* = *z × x*. **b-d** Nucleosome–nucleosome interaction configurations, *r* is the center-to-center distance between two nucleosomes.

Below, we explain the procedure used to categorize the relative orientation of the nucleosomes. We define the angles *{α, β*_*i*_, *β*_*j*_*}* as follows:

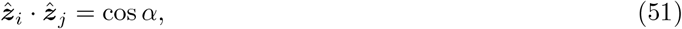

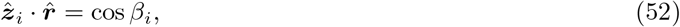

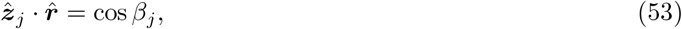

where 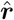 is the unit vector pointing from the center of the *i*^th^ nucleosome to the center of the *j*^th^ nucleosome, and 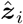 and 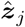 are the unit z-axis vectors of *i*^th^ and *j*^th^ nucleosomes respectively. We then use algorithm 1.

**Algorithm 1** Categorization of nucleosome-nucleosome orientation.

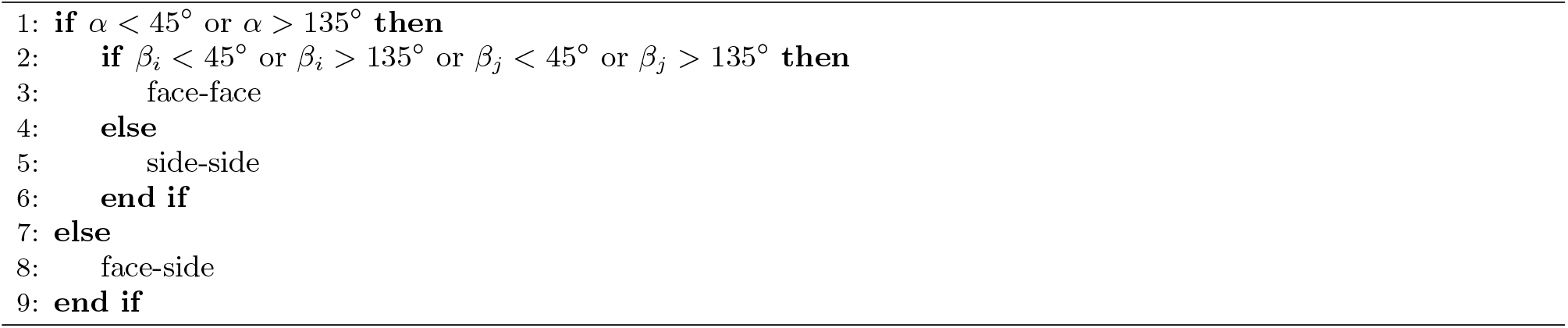

We then construct three interaction matrices 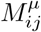 between the *i*^th^ and *j*^th^ nucleosomes, one for each relative orientation *µ* = *{*ff, fs, ss*}*:

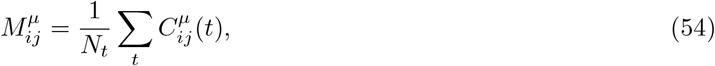

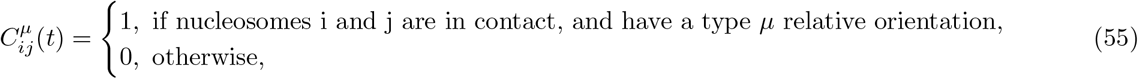

where *t* is the timestep. The sum is taken over all *N*_*t*_ snapshots used in the analysis. Two nucleosomes are defined to be in “contact” when the center to center distance between them is *<* 110 Å. The interaction matrices can then be projected onto a 1D map to describe the relative intensity of interactions between nucleosomes separated by (*k* − 1) neighbors,

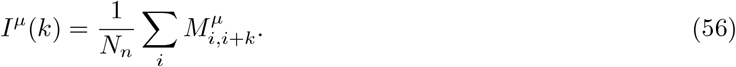

#### 2. Sedimentation coefficient

The sedimentation coefficient is computed using the HullRad method [42]:

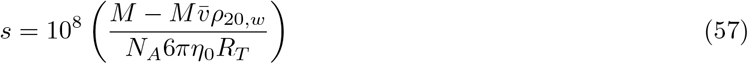

Where *M* is the total molar mass, 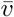 is the total partial specific volume, *ρ*_20,*w*_ is the density of water at 20C, and *η*_0_ is viscosity of water at 20C.

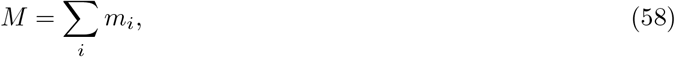

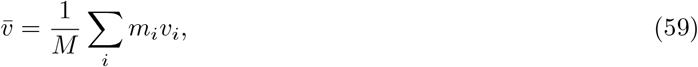

where *m*_*i*_ and *v*_*i*_ are the molar masses and specific volumes of the individual beads. *R*_*T*_ is the translational hydrodynamic radius which is computed via generating the convex-hull of the molecule. Full details of this calculation can be found in [42].

#### 3. mount of unwrapped DNA

For the simulations of breathing chromatin it becomes slightly difficult to define which DNA beads are nucleosomal and which are linker. This is because at the higher salt values the dense chromatin structures have DNA in contact with the nucleosome core proteins that is not part of that nucleosome, so simply computing the protein-DNA contacts will not work. To overcome this we developed the follow procedure: first we record which protein beads are bonded to the DNA in simulations of non-breathing nucleosomes, these protein beads are located circularly around the nucleosome in the locations where the DNA is typically wrapped. Then, for each frame in the breathing trajectory, we compute the contacts between the DNA and the aforementioned protein beads. For each nucleosome we now have a list of bound DNA beads. We then compute the median DNA bead in terms of index along the DNA sequence. This is approximately the center bead of that nucleosome’s nucleosomal DNA. We then look forwards and backwards along the DNA sequence, within the range of maximum and minimum indices of the bound DNA beads, and unless a large continuous region of unbound DNA (*>*100bp) is found, all the DNA between the maximum and minimum limits is added. Each nucleosome now has a contiguous section of nucleosomal DNA assigned to it. Finally the list of nucleosomal DNA is checked for overlaps and any are removed to ensure that each DNA bead can only be a member of one nucleosome. The average amount of unwrapped DNA per nucleosome is then computed as:

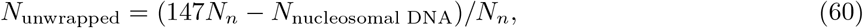

where *N*_*n*_ is the number of nucleosomes, *N*_nucleosmal DNA_ is the total number of nucleosomal DNA beads, and 147 is the typical number of base pairs of DNA wrapped round one nucleosome.

#### 4. Nucleosome valency

Nucleosome valency *V*, is defined as the average number of other nucleosomes a nucleosome is in contact with, as defined in Section VIII A 1.

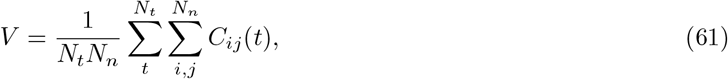

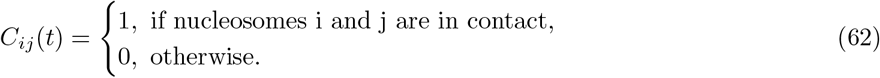

#### 5. Nucleosome–nucleosome contacts

To calculate the inter-nucleosome molecular contacts we use a similar procedure to the nucleosome-nucleosome interactions but at bead level rather than nucleosome level. For each timestep in the simulation trajectory the total contact matrix is computed for all beads.

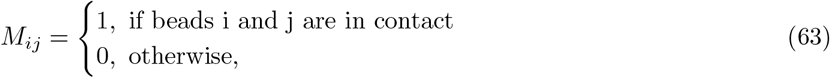

where “in contact” here is true when the distance between beads i and j is less than ((*σ*_*i*_ + *σ*_*j*_)*/*2 + 1Å). For each bead three sums are performed. One each counting up the contacts with: DNA, Histone tail, and Globular domain beads. Importantly the contacts are only counted if they are located in different nucleosomes. For the breathing DNA this is non-trivial. To proceed we defined nucleosomal DNA using the same method as in Section VIII A 3. The remaining linker DNA is then assigned to the nucleosome it is closest to (in term of DNA sequence, not spatial distance). This process enables the contacts to be computed for interactions between different nucleosomes that would otherwise be dominated by the intra-nucleosome contacts.

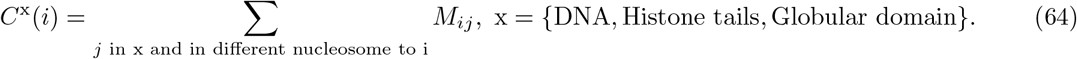

*C*^x^(*i*) is then averaged over all nucleosomes and all timesteps in the trajectory, and normalized by its maximum value *C*_MAX_. The values of *C*^x^(*i*) are plotted in Figure 4a and 4c. To generate the visualizations in Figure 4b and 4d each bead is given a RGB color according to:

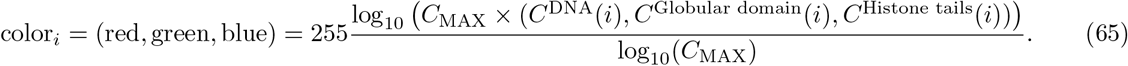

Where RBG values are integers in the range 0-255.

### B. Analysis of minimal coarse-grained simulations

#### 1. Radius of gyration

To enable comparison with the chemically-specific model the radius of gyration is computed using only the core beads:

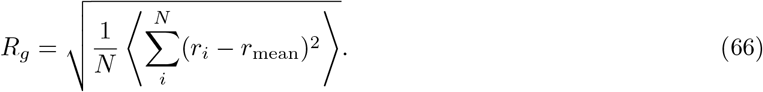

#### 2. Estimation of liquid-network connectivity

We define the connectivity as the mean number of connections per chromatin array in the high-density phase multiplied by the density of high-density phase, which gives the connectivity as the number of inter-chromatin bonds per unit volume. The number of connections of a chromatin array is defined as the number of distinct chromatin arrays it is in contact with. Two nucleosomes are defined to be “in contact” if the nucleosome–nucleosome distance (i.e. any nucleosome in one chromatin array relative to any nucleosome in another chromatin array), is less than 110Å.

## IX. ADDITIONAL RESULTS

Using our chemically-specific model, we performed additional simulations for 12-nucleosome arrays with uniform NRL of 195 bp (Figure S10).

**Figure S10.**
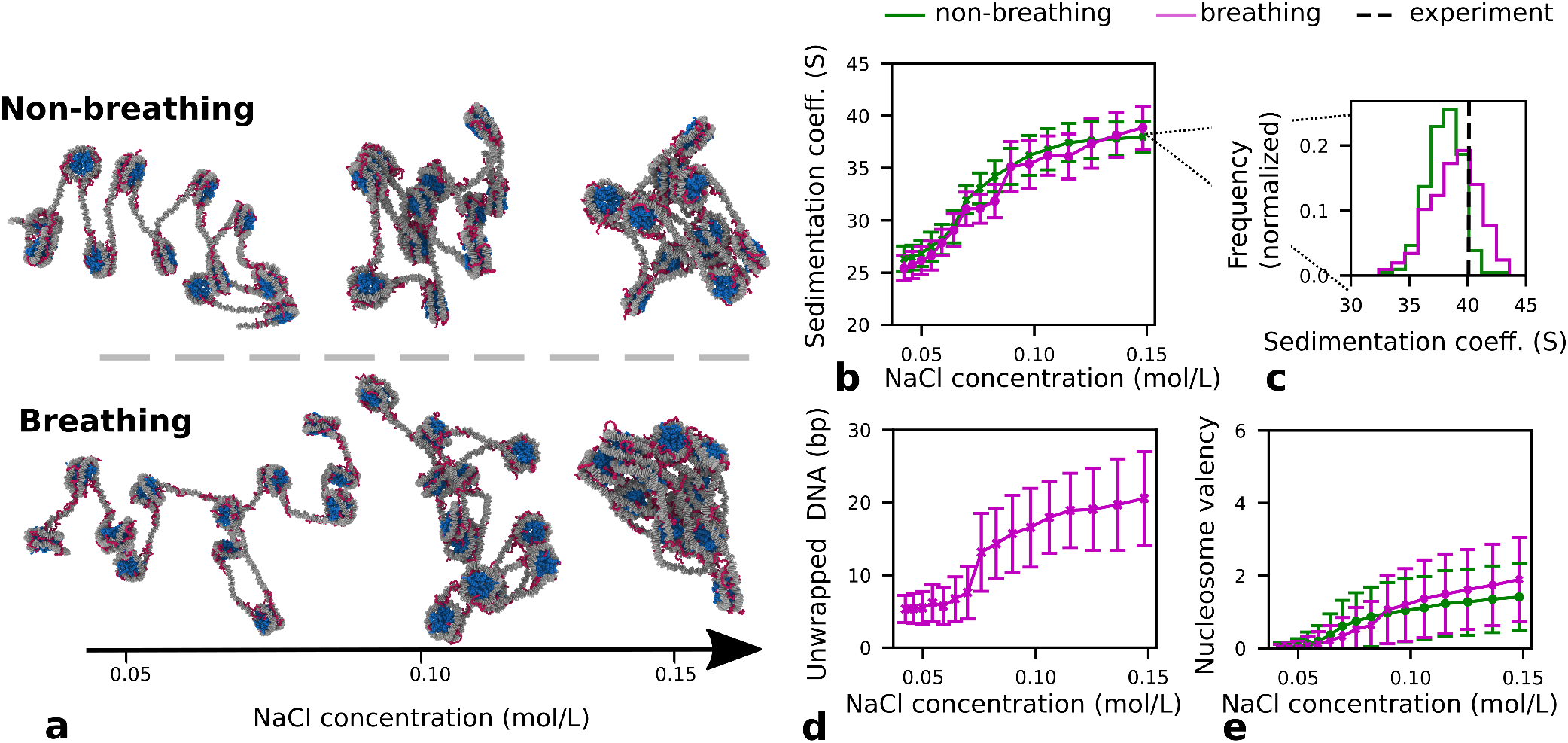
12-nucleosome 195 bp NRL chromatin simulations. **a** Representative simulation snapshots of 195-bp 12-nucleosome chromatin with non-breathing (top) versus breathing (bottom) nucleosomes at three different salt concentrations: 0.05 mol/L, 0.10 mol/L and 0.15 mol/L of NaCl. **b** Sedimentation coefficients versus NaCl concentration (right) for chromatin with non-breathing (green) versus breathing (magenta) nucleosomes. **c** Histograms comparing the distributions of sedimentation coefficient values for chromatin with non-breathing (green solid) and breathing (magenta solid) at 0.15 mol/L in our simulations with the experimental value from reference [43] (black dashed). **d** Number of average DNA bp that unwrap per nucleosome in our simulations at varying concentration of NaCl. **e** Average nucleosome valency versus concentration of NaCl.

